# Opposing Roles for the Spectraplakin Short Stop in Stable and Dynamic Dendrites Reveal Divergent DLK Signaling and a Role in Dendrite Regeneration

**DOI:** 10.64898/2026.09.06.749732

**Authors:** Vinicius N. Duarte, Rostislav Brichko, Viana Najafi, Katherine L. Thompson-Peer

## Abstract

Dendrites are vital to neuronal function, and dendrite injury occurs after neurological traumas such as stroke, traumatic brain injury, or neurodegenerative diseases. Despite their importance, the mechanisms underlying dendrite maintenance or regeneration remain poorly understood. The *Drosophila* gene *short stop* (*shot*), orthologous to ACF7/MACF1 in mammals, functions as an actin-microtubule crosslinker during neuronal development. Here, we investigate *shot’s* role in dendrite stability and repair using *Drosophila* sensory neurons. We find that *shot* plays opposing, cell type-specific roles in dendrite maintenance: it restricts excessive branch growth in simple, stable neurons, while being required for dendrite coverage in complex, dynamic neurons. These opposing functions are reflected in distinct localization patterns in stable versus dynamic dendrite arbors. Loss of *shot* destabilizes the microtubule cytoskeleton in stable neurons, and activates Wallenda/DLK signaling in both stable and dynamic neurons. Downstream of Wallenda/DLK, JNK signaling diverges between neuron types, with canonical basket/JNK activation occurring only in neurons with stable dendritic arbors. After injury, *shot* promotes dendrite regeneration in both neuron types and accumulates in distinct shapes in regenerated dendrites, with specific domains critical for proper Shot accumulation patterns. Collectively, these findings establish *shot* as a context-dependent regulator of dendrite maintenance and repair, and demonstrate that the structural identity of a dendritic arbor shapes how neurons sense and respond to cytoskeletal perturbation.

**Graphical Abstract:** 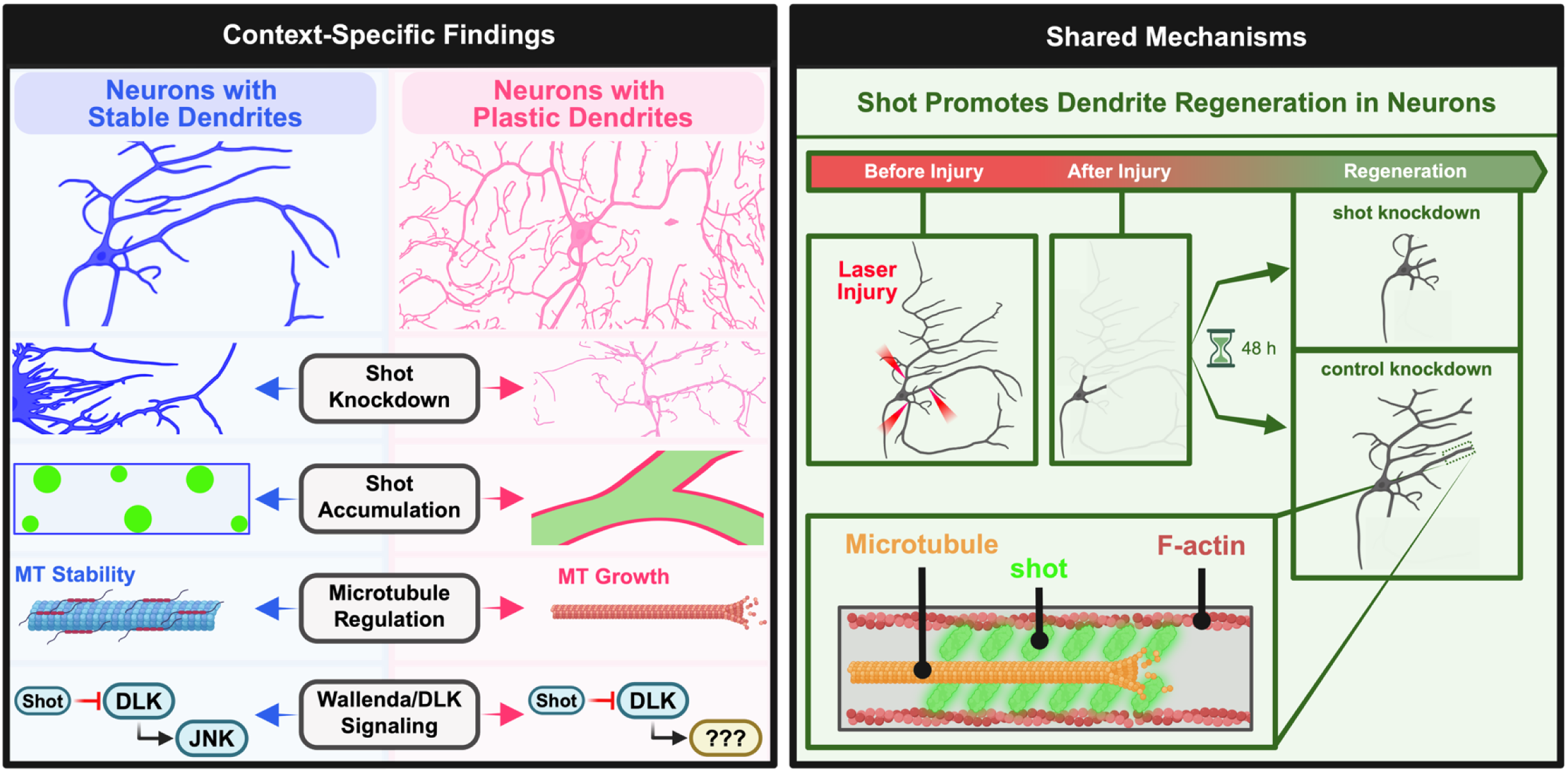

## Introduction

Dendrites are essential to neuronal function, acting as cellular antennae that receive signals from the environment or synaptic partners. Dendrite injury has been observed in human tissue following nervous system injuries such as traumatic brain injury (TBI) and stroke, and changes to dendritic architecture have been documented in aging and neurodegenerative disease. For example, levels of the somatodendritic marker MAP2 correlate with functional recovery after TBI in brain tissue biopsies from living patients, and MAP2 immunoreactivity is decreased in postmortem human tissue of individuals who suffered from ischemia/hypoxia relative to controls (Kühn et al., 2005; Yip et al., 2022). In the context of aging, progressive changes to dendrite integrity have been documented in the human prefrontal, motor, and temporal cortices, and early dendritic dystrophy has been observed in brain tissue from individuals with primary age-related tauopathy (Scheibel et al., 1975; Nakamura et al., 1985; Brabander et al., 1998; Shi et al., 2020). Thus, dendrite injury is clinically relevant across many neurological disorders. Investigating how dendrites are maintained or replaced after injury may therefore uncover novel therapeutic targets.

Early studies examining the responses to dendrite injury concluded that dendrite regeneration was not possible (Gross et al., 1983; Lucas et al., 1985; Emery et al., 1987; Gross and Higgins, 1987; Lucas, 1987; Sacconi et al., 2007). However, more recent studies in *Drosophila*, *C. elegans*, and *D. rerio* have established frameworks to study this biological process in live, intact organisms (Song et al., 2012; Oren-Suissa et al., 2017; Stone et al., 2022). From both fly and worm studies using sensory neurons of the peripheral nervous system (PNS), we have learned that dendrite regeneration operates independently of *Wallenda*/DLK, which mediates a key signaling cascade for the regeneration of axons (Stone et al., 2014; Brar et al., 2022). Specifically, Stone et al. (2014) showed that dendrite regeneration is not impaired in class IV ddaC neurons when *Wallenda*/DLK is genetically reduced, and that the canonical DLK/JNK activity reporter Puc::GFP is not activated following dendrite injury in class I ddaE neurons. This lack of change in Puc::GFP levels after dendrite injury contrasts the increase in Puc::GFP levels observed after axon injury in those same neurons, indicating that acute dendrite severing does not engage the canonical DLK/JNK cascade. Collectively, this and other work suggests that dendrite regeneration is robust, and that neurons may engage signaling pathways distinct from those involved in axon injury to sense and respond to dendrite injury.

We chose to investigate the *Drosophila* gene *short stop* (hereafter *shot*; orthologous to mammalian Microtubule-Actin Crosslinking Factor 1 or ACF7/MACF1) as it promotes developmental dendrite growth and inhibits DLK signaling in neurons (Prokop et al., 1998; Gao et al., 1999; Lee et al., 2000; Valakh et al., 2013). However, much of what is known regarding *shot*’s function in neurons comes from studies examining the axonal compartment. For example, Shot regulates axon extension by guiding growing microtubules during *de novo* development (Kolodziej et al., 1995; Lee et al., 2000; Sanchez-Soriano et al., 2009; Alves-Silva et al., 2012; Ka and Kim, 2016; Hahn et al., 2021). Shot also maintains cytoskeletal stability in axons by serving as an actin-microtubule crosslinker (Applewhite et al., 2010, 2013; Alves-Silva et al., 2012; Okenve-Ramos et al., 2024). After axon injury, *shot* mutants display accelerated axonal sprouting due to activation of the DLK signaling pathway (Valakh et al., 2013). In dendrites, our understanding of *shot*’s function is restricted to its roles in development. In *Drosophila* neurons, *shot* mutants display defects in dendrite morphogenesis (Prokop et al., 1998; Gao et al., 1999; Nithianandam and Chien, 2018), and some protein domains of Shot are dispensable for rescuing these morphological defects (Bottenberg et al., 2009). In mammals, ACF7/MACF1 regulates primary dendrite number, apical dendrite orientation, total dendrite length, and primary dendrite thickness of cortical pyramidal neurons (Ka and Kim, 2016). However, *shot*’s role in dendrite maintenance and regeneration are unknown.

In this study, we used an RNAi-knockdown approach to reduce *shot* expression after initial dendrite development. This allowed us to interrogate *shot’s* roles in dendrite maintenance and regeneration. We examined the consequences of *shot* knockdown in morphologically distinct dendritic arborization neurons of the *Drosophila* PNS (Grueber et al., 2002, 2003). These distinct morphologies are specified, in part, by differential cytoskeletal stability: class I ddaE neurons have stable dendrite arbors, while class IV ddaC neurons have plastic ones (Das et al., 2017; Nanda et al., 2023). We show that *shot* has opposing, cell-type specific roles: *shot* restricts excessive dendrite growth in stable neurons, but promotes higher-order branch growth in highly dynamic dendritic arbors. These distinct roles in dendrite growth are reflected in Shot’s unique localization patterns in stable versus dynamic arbors. We then show that *shot* knockdown lowers microtubule stability in simple, stable arbors, and that *shot* loss activates *Wallenda*/DLK signaling in both cell types, with canonical JNK signaling only activated in neurons with stable dendrite architecture. Lastly, we find that *shot* promotes dendrite regeneration after injury, and that full-length Shot accumulates in regenerating dendrites, with specific protein domains required for proper accumulation patterns.

## Results

### Shot loss alters dendrite morphology in a cell type-specific manner

We set out to identify a potential role for *shot* in dendrite outgrowth and maintenance. To achieve this, we used the 2.21-Gal4 (class I da neuron driver) and ppk-Gal4 (class IV da neuron driver) to knock down *shot* using RNAi (shot^HMJ23381^). Each of these drivers begins expression in late embryogenesis, after initial dendrite outgrowth (Grueber et al., 2003, 2007; Sugimura et al., 2003). Throughout this study, we used *yTub37C*.RNAi as a control as the *yTub37C* gene is not expressed in somatic tissues including neurons, which instead rely on *yTub23C* for microtubule nucleation (Wiese and Schmid, 2008; Nguyen et al., 2014). We imaged *yTub37C* knockdown neurons or *shot* knockdown neurons in late first-instar larvae at 48 hours after egg lay (h AEL) and late third-instar larvae at 144 h AEL. Class IV ddaC neurons with *shot* knockdown displayed reductions in territorial coverage at both 48 h AEL (Figure 1a-b) and 144 h AEL (Figure 1c-d). In class I ddaE neurons, *shot* knockdown resulted in no significant differences in branch number or total dendrite length at 48 h AEL, but these neurons displayed a significant increase in branch number at 144 h AEL (Figure 1e-f). Taken together, these data demonstrate that *shot* serves distinct functions in the stable, simple dendrite arbors of class I ddaE neurons versus the dynamic, complex dendrite arbors of class IV ddaC neurons. In class I neurons, *shot* inhibits excessive branching, while in class IV neurons, *shot* promotes dendrite branching. Moreover, plastic dendrite arbors may rely more heavily on *shot* function as defects in territorial coverage were detectable in class IV ddaC neurons as early as 48 h after egg laying (Figure 1a).

**Figure 1:**
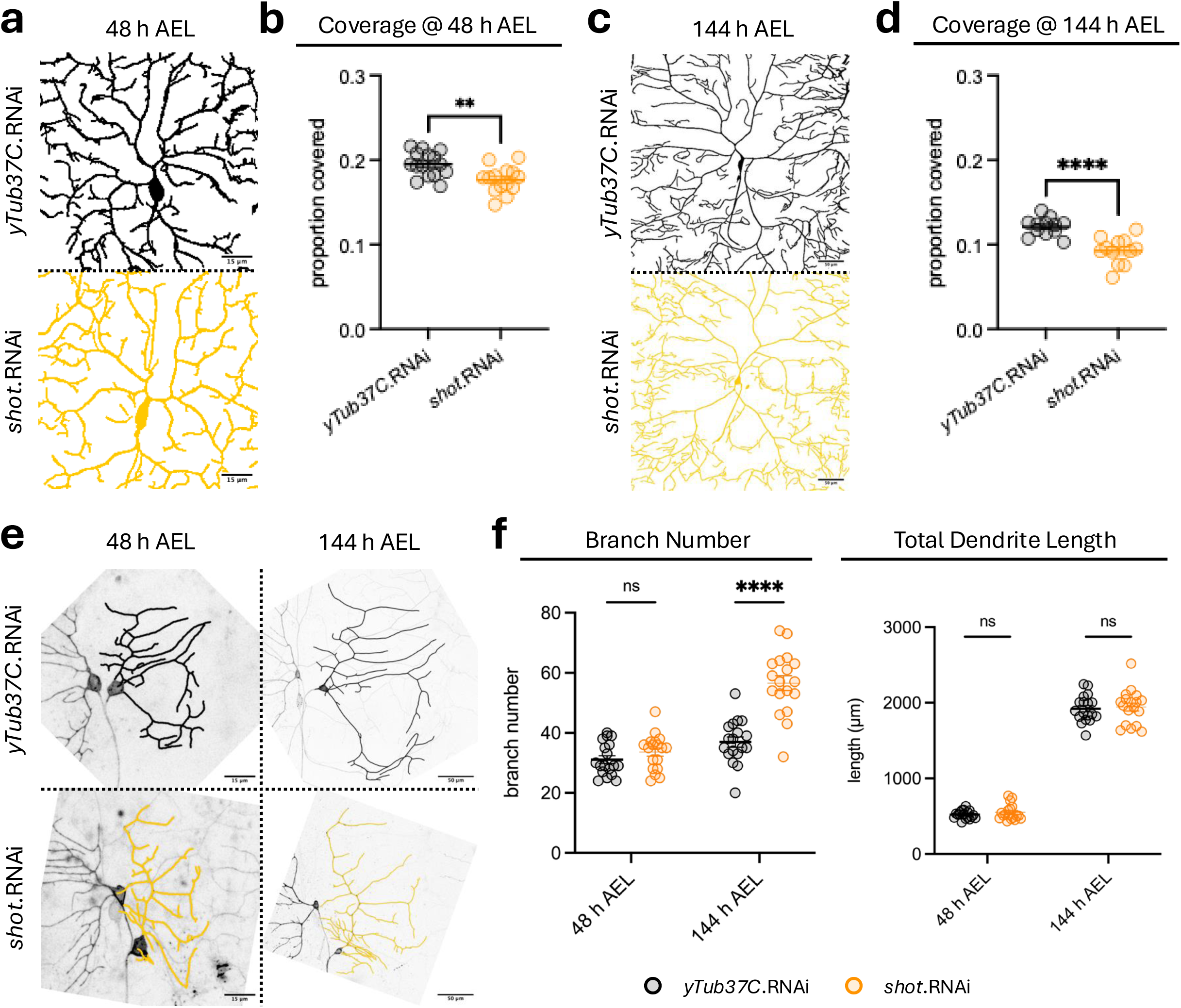
*Shot* knockdown alters dendrite arbor morphology in a developmental and cell-type specific manner. **(a)** Class IV ddaC neurons in 48 h AEL larvae with control *yTub37C* knockdown vs *shot* knockdown. **(b)** Territory coverage showed a significant decrease in neurons with *shot* knockdown. Welch’s t-test. p = 0.0024. N = 14-15 cells/group, 4 larvae/group, 3-4 cells/larvae. **(c)** Class IV ddaC neurons in 144 h AEL larvae with control *yTub37C* knockdown vs shot knockdown. **(d)** Territory coverage showed a significant decrease in neurons with shot knockdown. Welch’s t-test. p < 0.0001. N = 12 cells/group, 4 larvae/group, 3 cells/larvae. **(e)** Class I ddaE neurons tracked from 48 h AEL to 144 h AEL with control *yTub37C* knockdown vs *shot* knockdown. **(f)** Branch number showed a significant increase in neurons with *shot* knockdown at 144 h AEL. Total dendrite length resulted in no significant difference between groups at either developmental stage. Multiple unpaired t-tests with Welch correction, with Holm-Šídák multiple comparisons correction. Adjusted p < 0.000001 for branch number comparison at 144h AEL. N = 18 cells/group, 6 larvae/group, 3 cells/larvae. Mean ± SEM is plotted for all graphs.

To confirm that our *shot* knockdown approach was lowering Shot protein levels, we immunostained for Shot in late third-instar larval fillet preparations with Gal4[2.21] > UAS-CD4::tdTomato labeled class I ddaE neurons (Figure S1a). *Shot* knockdown resulted in significantly lower anti-Shot intensity in ddaE somas (Figure S1b), confirming effective RNAi-mediated reduction of Shot protein expression. We then tested the ability of a second *shot* RNAi line (*shot*^GLO1286^, hereafter referred to as *shot*.RNAi (B)) to recapitulate the maintenance phenotypes we identified with *shot*^HMJ23381^ in Figure 1. In class IV ddaC neurons, the data trended toward decreased territorial coverage (p = 0.0506) for *shot* knockdown neurons imaged at 144 h AEL (Figure S1c-d). However, *shot*.RNAi (B) had no significant effect on branch number or total dendrite length in class I ddaE neurons imaged at 48 h, 96 h, or 144 h AEL (Figure S1e-f).

Prior studies demonstrated that classical *shot* mutants disrupt dendrite development in class I ddaE neurons and class IV ddaC neurons (Gao et al., 1999; Nithianandam and Chien, 2018). We next sought to determine whether these developmental defects could be recapitulated with our *shot* RNAi knockdown approach using two distinct methods. First, we employed the 109(2)80-Gal4 driver that drives expression in all da neurons prior to initial dendrite outgrowth (Gao et al., 1999). Knocking down *shot* (*shot*^HMJ23381^) with this early pan-da neuron Gal4 driver resulted in significant decreases in territorial coverage for class IV ddaC neurons imaged at 144 h AEL (Figure S1g). Second, we assessed the effect of *shot* knockdown in adult class IV v’ada neurons, which prune all larval dendrites during pupariation and elaborate a new dendrite arbor in the adult fly abdomen (Kuo et al., 2005; Shimono et al., 2009). Our ppk-Gal4, UAS-CD4::tdTomato labeling approach resulted in both class IV v’ada neurons and class II neurons expressing CD4::tdTomato in adults (Figure S1h, purple arrows point to class IV v’ada somas and red arrows point to class II soma). We therefore restricted quantification to the region near the ventral midline, as this area is exclusively class IV v’ada dendrites and receives minimal contribution from class II neuron dendrites (Shimono et al., 2009). In these v’ada-only regions, *shot* knockdown resulted in significant decreases in territorial coverage of the adult abdomen imaged at 5 days post-eclosion (Figure S1h). Together, these results showed that early *shot* knockdown prior to initial dendrite outgrowth decreases territorial coverage in both larval and adult class IV da neurons. These cell type-specific effects on dendrite morphology prompted us to examine whether Shot itself is distributed differently across these two neuron types.

### Dendritic localization of shot is cell type-specific

Shot localization has primarily been studied in developing axons, where it localizes to growth cones to promote MT polymerization (Sanchez-Soriano et al., 2009; Alves-Silva et al., 2012; Hahn et al., 2021). In dendrites, Shot is recruited to growing dendrite tips in class IV ddaC neurons of first-instar larvae (Davies et al., 2025). Given *shot’s* roles in both MT extension and MT bundle formation, we assessed its localization in stable class I ddaE neurons and plastic class IV ddaC neurons in third-instar larvae imaged at 120 h AEL to capture more of the dendrite arbor in older larvae. To visualize Shot localization in these neurons, we expressed a GFP-tagged full-length Shot isoform (hereafter Shot.LA) using the 2.21-Gal4 and ppk-Gal4 drivers. Endogenous Shot is not reliably detectable in dendrites by immunohistochemistry due to dim signal (Figure S1a). In stable class I ddaE neurons, we observed small punctate Shot.LA accumulations throughout the dendrite arbor, and rod-like accumulations in the soma and axon (Figure 2a). The punctate localization of Shot.LA in dendrites is most obvious when the image is thresholded (Figure 2b). In dynamic class IV ddaC neurons, Shot.LA accumulated at dendritic branchpoints and was either diffuse or absent in dendritic shafts (Figures 2c-d). Somas contained high Shot.LA signals, and axons displayed rod-like accumulations (Figures 2c-d).

**Figure 2:**
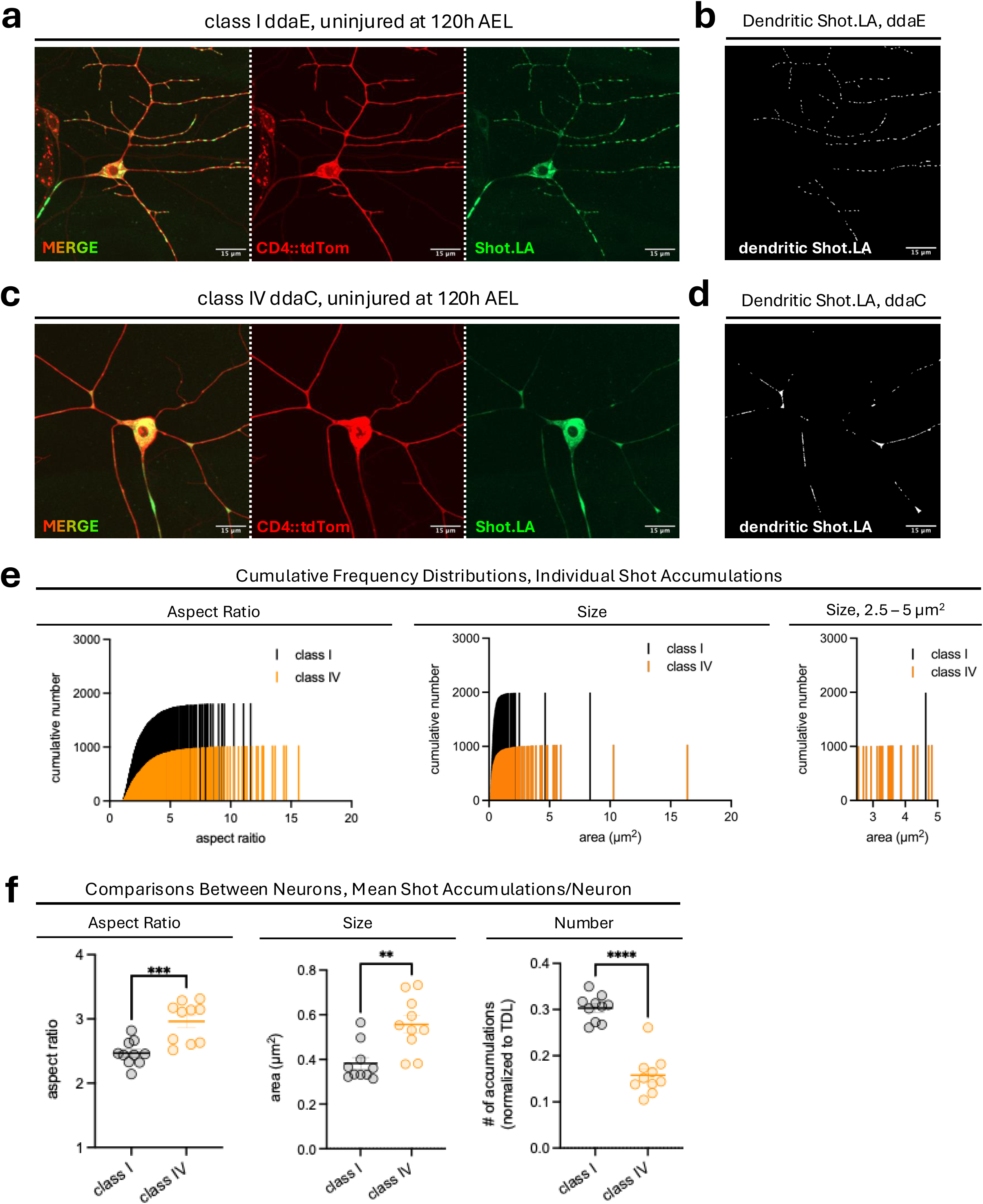
Shot exhibits distinct localization patterns in stable class I ddaE neurons versus dynamic class IV ddaC neurons. **(a)** Class I ddaE neurons expressing full-length, GFP-tagged Shot.LA at 120 h AEL. Neurons are labeled using UAS-CD4::tdTomato. **(b)** Thresholded Shot.LA signal only in class I ddaE dendrites, used for quantifications in (e) and (f). **(c)** Class IV ddaC neurons expressing full-length, GFP-tagged Shot.LA at 120 h AEL. Neurons are labeled using UAS-CD4::tdTomato. **(d)** Thresholded Shot.LA signal only in class IV ddaC dendrites, used for quantifications in (e) and (f). **(e)** Frequency distributions plotting aspect ratio or size of individual Shot.LA puncta in class I ddaE neurons or class IV ddaC neurons. Perfectly circular puncta are assigned an AR value of 1, whereas linear puncta are greater than 1. **(f)** Number, aspect ratio, and size of mean Shot.LA puncta per neuron in class I ddaE and class IV ddaC neurons. Welch’s t-test. Mean ± SEM. N = 10 cells/neuron type, 5 larvae/neuron type, 2 cells/larvae.

To quantify differences in Shot.LA protein localization between stable and plastic neurons, we plotted cumulative frequency distributions of aspect ratio (AR) and size for all individual Shot.LA accumulations in class I or class IV da neurons. An AR of 1 is perfectly round, while a higher AR indicates more oblong shape. The AR distribution was skewed towards round shapes for class I ddaE neurons, consistent with Shot.LA’s punctate accumulations in this stable neuron type, compared with more oblong shapes in class IV ddaC neurons (Figure 2e). Consistent with the AR frequency distribution, per neuron averages of Shot.LA accumulation ARs in class IV ddaC neurons were also significantly greater than those in class I ddaE neurons (Figure 2f). The size distribution was skewed left for both neuron classes, but class IV ddaC neurons had a cluster of larger accumulations roughly 2.5 - 5 µm^2^ in size (Figure 2e). These larger accumulations likely represent Shot.LA in dendritic branchpoints. Consistent with the size frequency distribution, per neuron averages of Shot.LA accumulation size in class IV ddaC neurons were significantly greater than those in class I ddaE neurons (Figure 2f). Lastly, the number of Shot.LA accumulations per unit total dendrite length (TDL) was significantly higher in class I ddaE neurons than in class IV ddaC neurons (Figure 2f). Taken together, we demonstrate that Shot accumulates differently in stable versus plastic dendrite arbors, with greater numbers of small, round accumulations in class I neurons, compared to fewer numbers of large, more oblong accumulations in class IV neurons. These distinct localization patterns suggest that Shot may serve different structural or regulatory roles in stable versus plastic dendrites, which we next explored by examining shot’s effects on the microtubule cytoskeleton.

### Shot knockdown destabilizes microtubules in stable dendrites

*Shot* promotes microtubule (MT) bundle formation via its MT-binding, C-terminal GAS2 domain (Lee and Kolodziej, 2002a). In *shot* mutant neurons, treatment with the MT-destabilizing agent nocodazole resulted in large gaps devoid of MTs that were not seen in wild-type neurons (Alves-Silva et al., 2012). Furthermore, axonal MT decay in aging neurons of the *Drosophila* visual system was exacerbated with *shot* knockdown. Expression of Shot’s C-terminal domain rescued these degenerative MT defects (Okenve-Ramos et al., 2024). We therefore determined whether *shot* knockdown affected MT stability in stable class I ddaE neurons and plastic class IV ddaC neurons. To assess MT stability, we used a Tau::GFP protein trap to visualize endogenous expression levels of the microtubule-associated protein Tau enriched in the sensory neurons of the larval body wall. Because Tau associates with microtubule shafts, Tau::GFP levels serve as a proxy for microtubule polymer content and stability. In class I ddaE neurons, we measured Tau::GFP levels in three compartments: the axon (a), the non-comb dendrite (nc), and the comb dendrite (c) (Figure 3a, white arrows). These compartments were analyzed separately given their significant differences in Tau::GFP levels in wild-type neurons (Figure S3a). *Shot* knockdown in these neurons resulted in significant decreases in Tau::GFP levels in the axon and comb dendrite, but not the non-comb dendrite (Figure 3a-b). In class IV ddaC neurons, we measured Tau::GFP levels in three compartments: the axon (a), primary dendrite branches (p), and terminal dendrite branches (t) (Figure 3c, white arrows). Terminal branches have minimal Tau::GFP and therefore served as a negative control. In these neurons, *shot* knockdown did not change Tau::GFP levels in any compartment (Figure 3c-d). Taken together, these data demonstrate that *shot* promotes MT stability in class I ddaE neurons but not class IV ddaC neurons, as assessed by Tau:: GFP levels.

**Figure 3:**
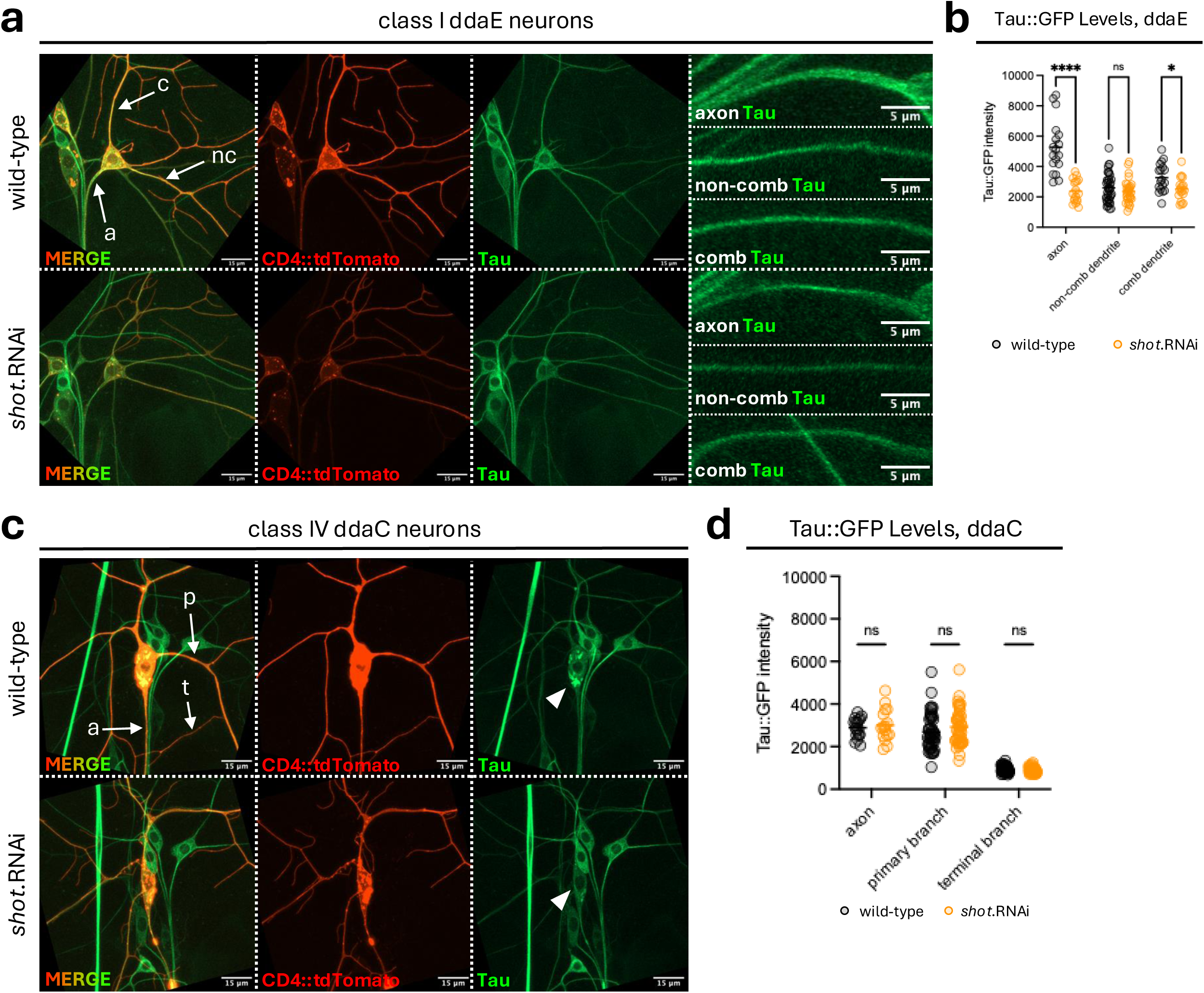
*Shot* loss decreases MT stability in class I ddaE but not class IV ddaC neurons. **(a)** Class I ddaE neurons in 144 h AEL larvae in wild-type vs *shot* knockdown. Neurons are labeled using 2.21-Gal4 and UAS-CD4::tdTomato. Stable microtubules are labeled using a Tau::GFP protein trap. **(b)** Tau levels showed a significant decrease in axons and comb dendrites, but not non-comb dendrites in neurons with *shot* knockdown. Multiple unpaired t-tests with Welch correction, with Holm-Šídák multiple comparisons correction. Adjusted p = 0.000005 for axon and 0.022999 for comb dendrite. N = 18 cells/genotype, 6 larvae/genotype, 3 cells/larvae, 18 axons/genotype, 18 comb dendrites/genotype, and 30-37 non-comb dendrites/genotype. **(c)** Class IV ddaC neurons in 144 h AEL larvae in wild-type or *shot* knockdown. Neurons are labeled using ppk-Gal4 and UAS-CD4::tdTomato. Stable microtubules are labeled using a Tau::GFP protein trap. **(d)** Tau levels showed no changes in neurons with *shot* knockdown. Multiple unpaired t-tests with Welch correction, with Holm-Šídák multiple comparisons correction. N = 16 cells/genotype, 4 larvae/genotype, 4 cells/larvae, 16 axons/genotype, 45-48 primary dendrites/genotype, and 48-51 terminal dendrites/genotype. Mean ± SEM for all.

In addition to promoting MT stability, *shot* promotes F-actin cytoskeletal stability via its actin-binding, N-terminal CH domains (Qu et al., 2022). Therefore, we sought to assess how *shot* knockdown affects actin stability in da neurons using GMA, an F-actin marker (Edwards et al., 1997). GMA levels were significantly decreased in the axon, non-comb dendrite, comb dendrite, and soma of class I ddaE neurons (Figure S3c-d). In class IV ddaC neurons, GMA levels remained unchanged in the axon but displayed significant reductions in the primary branches, terminal branches, and soma (Figure S3e-f). However, we noted that *shot* knockdown resulted in significant decreases in UAS-CD4::tdTomato expression (Figure S3b), indicating that *shot* knockdown reduces the transcriptional output of our 2.21-Gal4 and ppk-Gal4 drivers. Since GMA is also UAS-driven, observed changes in GMA levels could reflect reduced driver output rather than genuine alterations in F-actin stability, complicating interpretation of results with this approach. The role of *shot* in F-actin stability may be more reliably assessed using a Gal4/UAS-independent, neuron-specific protein trap marker that is not susceptible to variability in Gal4 driver output. The reduction in MT stability observed in class I ddaE neurons following *shot* knockdown prompted us to examine whether this cytoskeletal perturbation activates downstream kinase signaling.

### Shot loss triggers canonical and non-canonical DLK signaling

*Shot* stabilizes the axonal cytoskeleton (Alves-Silva et al., 2012), which inhibits dual leucine zipper kinase DLK (or *Wallenda* in *Drosophila*) signaling. For example, overgrowth of axon terminals caused by *shot* loss is suppressed by concomitant loss of *Wallenda (Wnd)* (Valakh et al., 2013). Thus, we determined whether the dendritic phenotypes observed with *shot* loss could be suppressed by concomitant *Wnd* knockdown. To achieve this, we knocked down either *yTub37C* (control), *shot* alone, *Wnd* alone, or *shot* and *Wnd* together in class I or class IV da neurons (Figure 4a). Loss of *shot* produced the dendrite branch increase (class I) or decrease (class IV) phenotypes as observed previously (Figure 1), while concomitant knockdown of *shot* and *Wnd* eliminated these phenotypes (Figure 4b). In class IV da neurons, we identified an increase in territory coverage in *Wnd* knockdown alone, a phenotype that was not previously observed with classical *Wnd* mutants (Wang et al., 2013). Therefore, *shot* knockdown regulates dendrite branching via *Wallenda*/DLK signaling regardless of whether dendrites are stable or dynamic.

**Figure 4:**
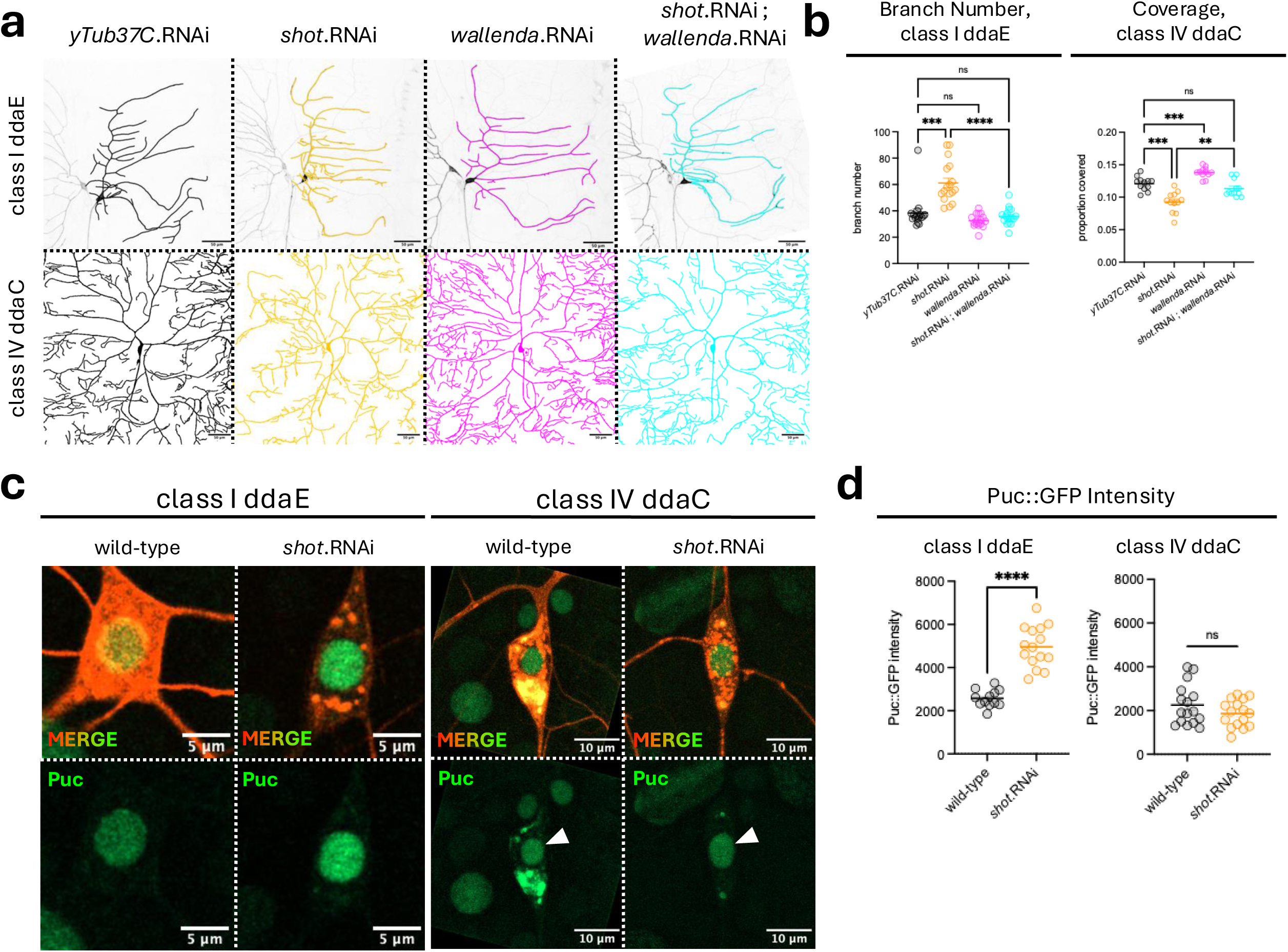
*Shot* knockdown activates DLK signaling in da neurons, but not canonical DLK/JNK signaling in class IV da neurons. **(a)** Class I ddaE (top row) and class IV ddaC (bottom row) neurons in 144 h AEL larvae with control *yTub37C*.RNAi, *shot*.RNAi, *Wallenda*.RNAi, or *shot*.RNAi and *Wallenda*.RNAi. **(b)** Left graph with branch number showed a significant increase in neurons with *shot* knockdown, and this was suppressed by simultaneous *shot* and *Wallenda* knockdown. Knockdown of *Wallenda* alone had no effect on dendrite branch number. Brown-Forsythe and Welch ANOVA tests with Dunnett T3 multiple comparisons corrections. N = 17 cells/genotype, 8 larvae/genotype, 2-3 cells/larvae. Right graph with territory coverage showed a significant decrease in neurons with *shot* knockdown, and this was rescued by simultaneous *shot* and *Wallenda* knockdown. Knockdown of *Wallenda* alone significantly increased territorial coverage. Brown-Forsythe and Welch ANOVA tests with Dunnett T3 multiple comparisons corrections. N = 12 cells/genotype, 4 larvae/genotype, 3 cells/larvae. **(c)** Puc::GFP in class I ddaE neurons and class IV ddaC in wild-type or *shot* knockdown neurons. Class I ddaE neurons are labeled using 2.21-Gal4 and UAS-CD4::tdTomato and class IV ddaC neurons are labeled using ppk-Gal4 and UAS-CD4::tdTomato. **(d)** Puc::GFP intensity in nuclei showed a significant increase in class I ddaE neurons with *shot* knockdown but no changes in Puc::GFP intensity were detected in class IV ddaC neurons with *shot* knockdown. Welch’s t-test. N = 12-15 cells/genotype, 4-5 larvae/genotype, 3 cells/larvae. Mean ± SEM is plotted for all graphs.

Canonical Wallenda/DLK signaling activates JNK, which phosphorylates c-Jun; c-Jun then partners with c-Fos to drive stress response gene transcription (Collins et al., 2006; Xiong et al., 2010). One JNK target gene, the dual-specificity phosphatase 10 (DUSP10; *puckered* or *puc* in flies), can be used as a reporter for JNK activity, and participates in a negative feedback loop for canonical DLK/JNK signaling (Martín-Blanco et al., 1998; Theodosiou et al., 1999). *Shot* knockdown increases levels of a Puc::LacZ enhancer trap in motor neurons of the larval ventral nerve cord, a phenotype that was again reversed with concomitant *Wnd* knockdown (Valakh et al., 2013). A GFP protein trap for *puc* has been used as a reporter of DLK/JNK activity in live, intact *Drosophila* larvae (Morin et al., 2001; Stone et al., 2014). Given that *shot* loss phenotypes were suppressed by concomitant *Wnd* knockdown in da neurons (Figure 4a-b), we next determined whether *shot* loss activates canonical *Wnd* signaling by measuring Puc::GFP levels (Figure 4c). We found a 2-fold increase in Puc::GFP levels in class I da neurons with *shot* knockdown, but no change in class IV da neurons (Figure 4d). Thus, *shot* loss activates canonical DLK/JNK signaling in stable class I da neurons, but likely activates non-canonical, JNK-independent DLK signaling in dynamic class IV da neurons. This non-canonical signaling could act through effectors like p38, which has been previously shown to mediate JNK-independent DLK signaling to regulate synaptic structure in fly motor neuron terminals (Klinedinst et al., 2013).

*Shot* loss phenotypes are suppressed by *Wnd* knockdown (Figure 4a-b), and *Wnd* overexpression is sufficient to activate DLK/JNK signaling (Collins et al., 2006). Based on these findings, we reasoned that *Wnd* overexpression should produce similar phenotypes to *shot* loss. Previous work has shown that *Wnd* overexpression does not change dendrite morphology of class I da neurons and significantly reduces the complexity of class IV da neurons (Wang et al., 2013). We reproduced these findings in class IV da neurons, as we found that *Wnd* overexpression phenocopied the reduction in dendrite coverage observed with *shot* loss, and indeed resulted in dendrite loss (Figures 5a and 5c). In contrast, *Wnd* overexpression failed to phenocopy the dendrite branch gain caused by *shot* loss in class I da neurons (Figure 5a-b).

**Figure 5:**
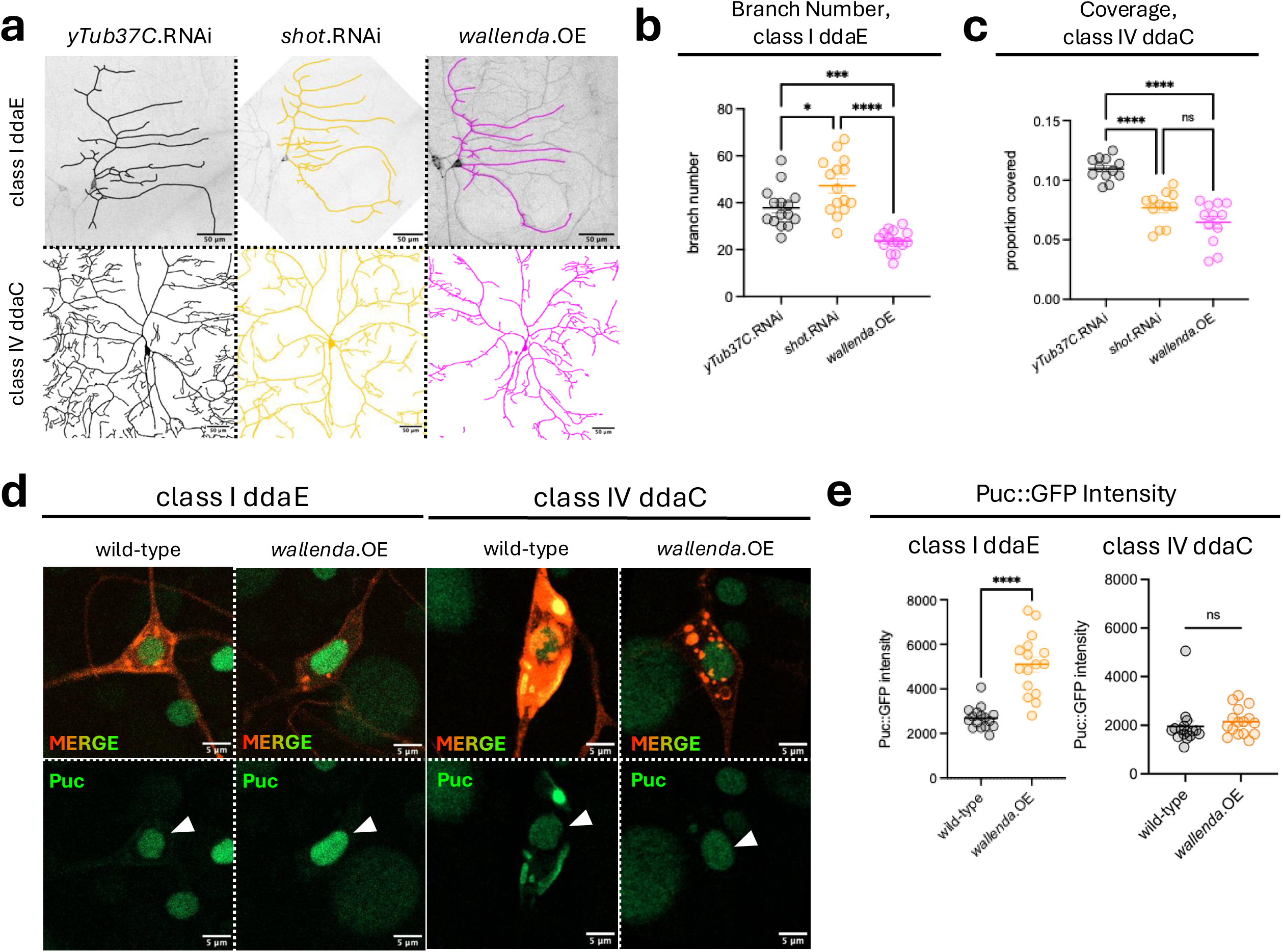
Wallenda/DLK overexpression phenocopies shot knockdown in ddaC but not ddaE neurons. **(a)** Class I ddaE (top row) and class IV ddaC (bottom row) neurons in 144 h AEL larvae with control *yTub37C*.RNAi, *shot*.RNAi, or *Wallenda*.OE. **(b)** Branch number showed a significant increase in neurons with *shot* knockdown, and this was not phenocopied by *Wallenda* overexpression. Ordinary one-way ANOVA with Tukey’s multiple comparisons corrections. N = 15 cells/genotype, 4 larvae/genotype, 3-4 cells/larvae. **(c)** Territory coverage showed a significant decrease in neurons with *shot* knockdown, and this was phenocopied by *Wallenda* overexpression. Ordinary one-way ANOVA with Tukey’s multiple comparisons corrections. N = 12 cells/genotype, 4 larvae/genotype, 3 cells/larvae. **(d)** Puc::GFP in class I ddaE neurons and class IV ddaC in wild-type larvae or in larvae overexpressing *Wallenda* with 2.21-Gal4 or ppk-Gal4. Class I ddaE neurons are labeled using 2.21-Gal4 and UAS-CD4::tdTomato and class IV ddaC neurons are labeled using ppk-Gal4 and UAS-CD4::tdTomato. **(e)** Puc::GFP intensity in nuclei showed a significant increase in class I ddaE and class IV ddaC neurons with *Wallenda* overexpression. Welch’s t-test. N = 16 cells/genotype, 4 larvae/genotype, 4 cells/larvae for both classes of neurons. Mean ± SEM is plotted for all graphs.

To validate that *Wnd* overexpression increases DLK/JNK signaling in da neurons, we measured Puc::GFP levels in class I or class IV da neurons overexpressing *Wnd* (Figure 5d). *Wnd* overexpression increased Puc::GFP levels in class I da neurons, but not class IV da neurons (Figure 5e). Notably, absence of change in class IV da neurons overexpressing *Wnd* mirrors the lack of a Puc::GFP change with *shot* knockdown (Figure 4d), providing independent evidence that class IV da neurons cannot activate canonical JNK signaling even when Wnd activity is directly elevated. Together with *shot* knockdown phenotypes being rescued by concomitant *Wnd* knockdown, these results suggest that *Wnd* activation may be necessary but insufficient to cause excessive branch growth in stable class I da neurons. In class IV da neurons, *Wnd* activity may be more tightly regulated by the upstream E3 ubiquitin ligase *highwire,* as *highwire* mutants did increase Puc::LacZ levels in these neurons (Wang et al., 2013). Having established that shot loss activates DLK signaling in both neuron types, we next examined whether shot also plays a role after acute dendrite injury.

### Shot promotes dendrite regeneration by localizing to regenerating dendrite tips

Loss of *shot* promotes axonal sprouting by mimicking a preconditioning “injury” via cytoskeletal destabilization and DLK/JNK activation prior to the insult itself (Valakh et al., 2013). In later stages of axon regeneration, *shot* likely promotes MT polymerization by acting as a MT +TIP (plus tip-interacting protein; Alves-Silva et al., 2012; Hahn et al., 2021). Given that regenerating dendrites display plus-end out MT growth (Song et al., 2012), we hypothesized that *shot* may promote dendrite regeneration. To test this, we severed all dendrite branches (complete dendrotomy) of control neurons or *shot* knockdown neurons in 96 h AEL larvae and assessed regenerative outcomes 48 h later, immediately prior to pupariation (Figure 6a). We chose to conduct injuries at 96 h AEL, as *Pten/Akt* signaling is dampened in da neurons at this time (Parrish et al., 2009), which allowed us to assess the role of *shot* in dendrite regeneration while minimizing developmental confounds. In class I ddaE neurons, *shot* knockdown resulted in significantly fewer regenerated dendrite branches and shorter total dendrite length (Figure 6b). This contrasted with the gain in dendrite branches we previously observed in uninjured *shot* knockdown neurons (Figures 1e and 1f). Using our second *shot* RNAi line, we found no significant changes to regenerated branch number or total dendrite length (Figure S6a-b); this absence of effect likely reflects its lower knockdown efficiency, consistent with the absence of a phenotype in uninjured class I ddaE neurons with this shot.RNAi(B) line (Figures S1e-f). *Shot* knockdown also impaired regenerative outcomes in class IV ddaC neurons, evidenced by significant reductions in regenerated dendrite area for both *shot* RNAi lines (Figure 6c-d and Figure S6c-d). Collectively, these data highlight the context-dependent functions of *shot:* while it displays cell type-specific roles in dendrite maintenance of uninjured neurons depending on cell type (Figure 1), it positively regulates dendrite regeneration in both cell types.

**Figure 6:**
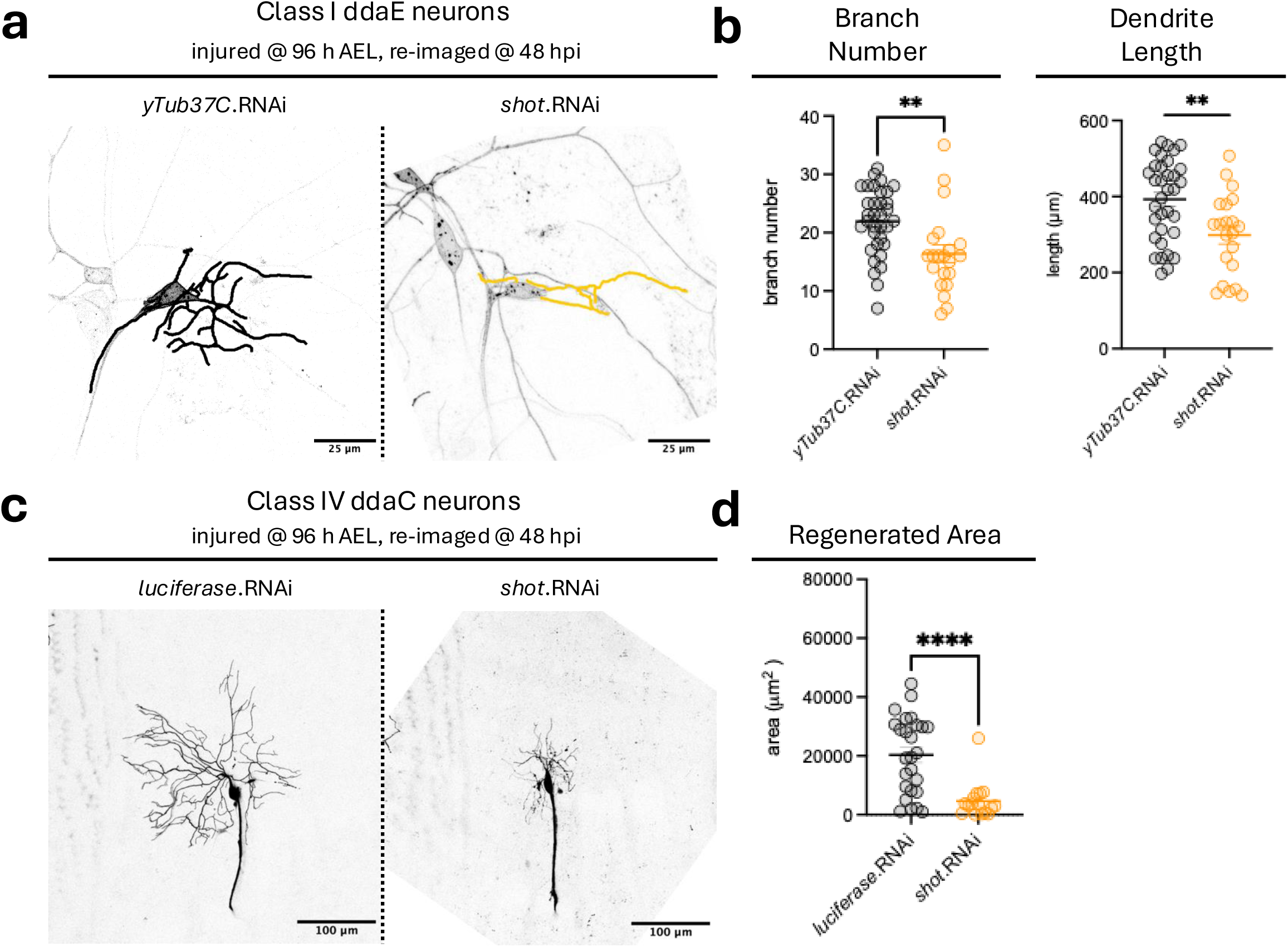
Shot knockdown impairs dendrite regeneration in class I ddaE and class IV ddaC neurons. **(a)** Class I ddaE neurons at 48 hpi in larvae with control *yTub37C* knockdown or *shot* knockdown. Injuries occurred 96 h AEL. Traces of regenerated dendrites are overlaid. **(b)** Branch number and total dendrite length showed a significant decrease in neurons with *shot* knockdown. Welch’s t-test. N = 21-34 cells/genotype, 10-12 larvae/genotype, 1-4 cells/larvae. **(c)** Class IV ddaC neurons at 48 hpi in larvae with control *luciferase* knockdown or *shot* knockdown. Injuries occurred 96 h AEL. Regenerated areas of dendrites are overlaid. **(d)** Regenerated area was significantly lower in neurons with *shot* knockdown. Welch’s t-test. N = 16-26 cells/genotype, 6-11 larvae/genotype, 2-3 cells/larvae. Mean ± SEM is plotted for all graphs.

*Drosophila shot* and mouse ACF7/MACF1 are essential for developmental neurite extension, displaying enriched localization at axonal growth cones and dendrite tips (Sanchez-Soriano et al., 2009; Ka and Kim, 2016). To test whether *shot* promotes dendrite regeneration by localizing to regenerating dendrite tips, we overexpressed GFP-tagged *shot* constructs in class I da neurons, severed all dendrite branches (complete dendrotomy) at 96 h AEL, and re-imaged neurons 48 h later (Figure 7). The full-length Shot isoform (Shot.LA, which was used to characterize Shot localization in uninjured dendrites in Figure 2) is present in the nervous system of *Drosophila*. We used a non-neuronal isoform of Shot (hereafter Shot.LC) as a comparison (Lee et al., 2000). Unlike full-length Shot.LA, Shot.LC contains just one of two N-terminal actin-binding domains and fails to rescue dendrite defects in *shot* mutants (Bottenberg et al., 2009). In regenerated dendrites, we observed rod-like accumulations of Shot.LA in regenerating dendrite tips (Figure 7a, arrows) and punctate Shot.LA accumulations throughout the entire dendritic arbor. However, Shot.LC only minimally localized to regenerating dendrites (Figure 7a). To quantify differences between the two isoforms, we measured Shot accumulation number per unit of total dendritic length (TDL) and Shot accumulation size in all regenerated dendrites. Dendritic Shot.LC accumulations were significantly fewer in number and smaller than Shot.LA accumulations (Figure 7b). Shot.LA accumulations were also present in regenerated class IV ddaC neuron dendrites, while Shot.LC accumulations were largely absent (Figure S7a).

**Figure 7:**
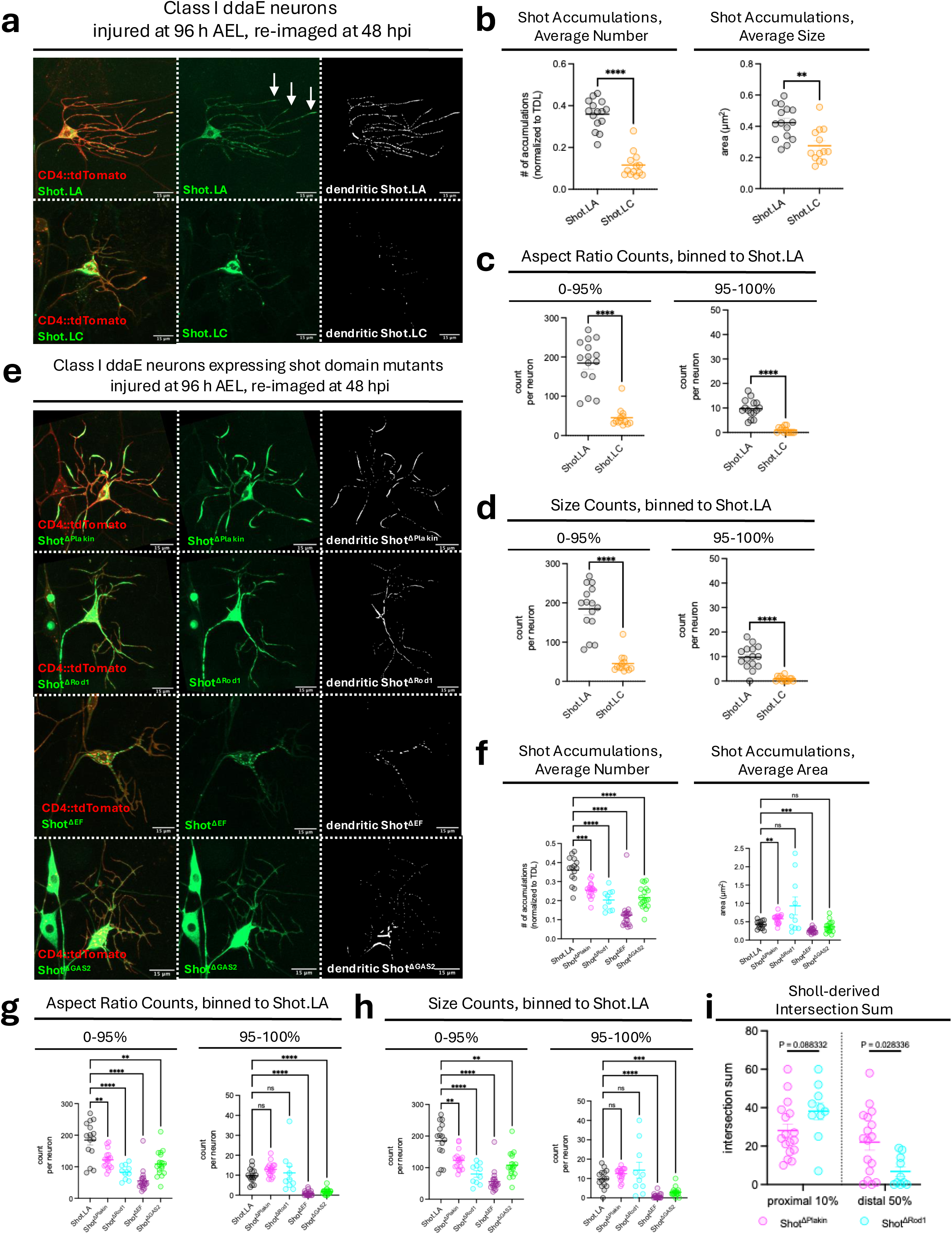
Shot domains distinctly regulate accumulations in regenerated dendrites. **(a)** Class I ddaE neurons at 48 hpi in larvae overexpressing GFP-tagged neuronal Shot.LA or non-neuronal Shot.LC. Injuries occurred 96 h AEL. Arrows point to rod-like Shot.LA accumulations in regenerating dendrite tips. **(b)** Shot puncta number, and Shot puncta size showed a significant decrease for both metrics in neurons expressing Shot.LC. **(c)** Per neuron counts of more punctate (aspect ratio = bin 0-95%) or more rod-like (aspect ratio = bin 95-100%) accumulations were significantly lower in Shot.LC relative to Shot.LA. **(d)** Per neuron counts of smaller (area = bin 0-95%) or larger (area = bin 95-100%) accumulations were significantly lower in Shot.LC relative to Shot.LA. Welch’s t-test. N = 13-15 cells/genotype, 3-5 larvae/genotype, 2-5 cells/larvae for a-d. **(e)** Class I ddaE neurons at 48 hpi in larvae overexpressing GFP-tagged Shot domain mutants. Injuries occurred 96 h AEL. **(f)** All Shot domain mutants showed a significant decrease in the number of accumulations (left graph). Shot accumulation size was significantly larger in neurons expressing Shot^ΔPlakin^ and significantly lower in neurons expressing Shot^ΔEF^ (right graph). **(g)** Per neuron counts of more punctate (aspect ratio = bin 0-95%) accumulations were significantly lower in all Shot domain mutants, while per neuron counts of more rod-like (aspect ratio = bin 95-100%) accumulations showed no change for Shot^ΔPlakin^ and Shot^ΔRod1^ relative to Shot.LA. **(h)** Per neuron counts of smaller (area = bin 0-95%) accumulations were significantly lower in all Shot domain mutants, while per neuron counts of larger (area = bin 95-100%) accumulations showed no change for Shot^ΔPlakin^ and Shot^ΔRod1^ relative to Shot.LA. Brown-Forsythe and Welch ANOVA with Dunnett’s T3 multiple comparisons correction was used for f-h. **(i)** Sholl-derived intersection sum in the proximal 10% showed a trending increase in intersection sum for Shot^ΔRod1^, while the distal 50% showed a significant decrease in intersection sum for Shot^ΔRod1^. Multiple unpaired t-tests with Welch correction, with Holm-Šídák multiple comparisons correction. N = 10-20 cells/genotype, 3-5 larvae/genotype, 2-5 cells/larvae for f-i. Mean ± SEM is plotted for all graphs. Bins in c, d, g, and h were established using control Shot.LA accumulations.

To differentiate between the punctate versus rod-like accumulations present in regenerated dendrites, the aspect ratio (AR) or size of accumulations were binned based on the Shot.LA measurements, as either punctate (0th–95th percentile) or rod-like (95th– 100th percentile). Per neuron counts of accumulation AR revealed significantly fewer punctate and fewer rod-like accumulations of Shot.LC relative to Shot.LA (Figure 7c). Similarly, per neuron counts of accumulation size revealed significantly fewer small and fewer large accumulations for Shot.LC relative to Shot.LA (Figure 7d). Taken together, these data suggest that full-length Shot promotes dendrite regeneration by facilitating microtubule extension at regenerating dendrite tips and by stabilizing regenerated dendrites throughout the arbor. Additionally, proper accumulation patterns in regenerated dendrites require both N-terminal actin-binding domains of Shot.

Shot’s central plakin and rod domains are dispensable for dendritic growth of embryonic motor neurons (Bottenberg et al., 2009). Its C-terminal Ca^2+^-binding domains regulate conformational states that govern Shot’s actin/microtubule crosslinking activity (Applewhite et al., 2013). In developing axons, Shot relies on its N-terminal actin-binding and C-terminal microtubule-binding domains to guide MT polymerization (Hahn et al., 2021). Thus, we examined which domains of Shot are necessary for proper accumulation morphology in regenerated dendrites. To do this, we overexpressed four GFP-tagged *shot* constructs lacking the plakin (Shot^ΔPlakin^), rod (Shot^ΔRod1^), Ca^2+^-binding (Shot^ΔEF^), or microtubule-binding (Shot^ΔGAS2^) domain. We performed complete dendrotomy at 96 h AEL and re-imaged neurons 48 h later (Figure 7e). We again measured Shot accumulation number per unit of total dendrite length and Shot accumulation size in all regenerated dendrites. Dendritic Shot accumulations for all domain mutants were significantly fewer in number than full-length Shot.LA accumulations (Figure 7f). With respect to size, Shot^ΔPlakin^ accumulations were significantly larger than Shot.LA accumulations, while Shot^ΔEF^ accumulations were significantly smaller (Figure 7f). When differentiating between the punctate and rod-like accumulations, there were significantly fewer punctate accumulations across all domain mutants but no change in rod-like accumulations for Shot^ΔPlakin^ or Shot^ΔRod1^ mutants (Figure 7g). Likewise, per neuron counts of accumulation size revealed significantly fewer small accumulations for all domain mutants but no changes in the number of large accumulations for Shot^ΔPlakin^ or Shot^ΔRod1^ mutants relative to Shot.LA (Figure 7h).

Despite Shot^ΔPlakin^ and Shot^ΔRod1^ mutants’ ability to accumulate in large, rod-like shapes, their localization phenotypes differed drastically (Figure 7e). This suggested distinct functional roles for these domains in regenerated dendrites. To quantify these potential differences in localization, we generated soma-centered Sholl plots using the dendritic Shot^ΔPlakin^ and Shot^ΔRod1^ accumulation masks (Sholl, 1953; Ferreira et al., 2014; Arshadi et al., 2021). The resulting Sholl plots detected more shot accumulations near the soma for Shot^ΔRod1^ mutants (proximal 10%), while Shot^ΔPlakin^ mutants had more accumulations farther away from the soma (distal 50%) (Figure 7i). This demonstrates that while both Shot^ΔPlakin^ and Shot^ΔRod1^ accumulate in large patches in regenerated dendrites, Shot^ΔPlakin^ mutants retain the ability to localize to regenerated dendrite tips, while Shot^ΔRod1^ has lost the ability to localize to dendrite tips. Collectively, these data obtained from *shot* domain mutants suggest that all domains of the Shot protein play a critical role in the resumption of punctate Shot accumulations in regenerated dendrites, that the plakin and rod domains of Shot are dispensable for its accumulation into a rod-like morphology, and that the plakin domain of Shot is dispensable for Shot’s ability to traffic to regenerating dendrite tips, suggesting that the plakin domain is not required for microtubule extension. These results establish shot as a positive regulator of dendrite regeneration and reveal distinct structural requirements for Shot’s accumulation and function in regenerating dendrites

## Discussion

Our findings establish shot as a context-dependent regulator of dendrite maintenance and repair, revealing that the cytoskeletal identity of a dendritic arbor fundamentally shapes how neurons interpret and respond to cytoskeletal perturbation We found that *shot* regulates dendrite arbor maintenance in a cell type-specific manner, restricting excessive dendrite branching in stable neurons and promoting dendrite growth in plastic neurons. We further showed that Shot accumulates in distinct manners between the two neuron types, suggesting different functional roles of *shot* in stable versus plastic dendrites. We found that *shot* promotes microtubule stability in stable neurons, but did not detect changes in microtubule stability in plastic neurons. We demonstrated that *shot* inhibits canonical DLK signaling via the JNK pathway in stable neurons, but *shot* inhibits DLK signaling in a JNK-independent manner in plastic neurons. Finally, we showed that *shot* promotes dendrite regeneration after injury in both neuron types, and all Shot protein domains are required for proper Shot accumulation in regenerated dendrites. Together, these findings advance our understanding of how the structural identity of a dendritic arbor governs the molecular pathways neurons use to maintain and repair their dendrites.

### Distinct Functions of Shot in Dendrite Maintenance and Development

The morphological consequences of *shot* loss have been studied in dendrites using various models including *Drosophila* CNS/PNS neurons and mouse cortical neurons *in vitro* and *in vivo*. Across all models, *shot* loss has consistently resulted in reductions to dendritic complexity (Prokop et al., 1998; Gao et al., 1999; Ka and Kim, 2016; Davies et al., 2025). This reduction in dendritic complexity is consistent with our findings that *shot* knockdown in plastic class IV da neurons resulted in decreased territorial coverage (Figure 1a-d). However, *shot* loss in class I da neurons increased dendrite branch number (Figure 1e-f), suggesting distinct context-dependent roles for *shot* in dendrite maintenance versus dendrite development. The idea that *shot* serves mechanistically distinct functional roles in dendrite maintenance is supported by the accumulation patterns of GFP-tagged Shot in both class I and class IV da neurons. As opposed to rod-like accumulations at growing neurite tips, Shot accumulated as small puncta throughout the arbor in class I da neurons and was diffuse and enriched at dendritic branchpoints in class IV da neurons (Figure 2). Furthermore, *shot’s* function in stabilizing dendritic microtubules in class I da neurons, but not class IV da neurons, provided additional evidence of context-dependent roles in dendrite maintenance.

*Shot* loss resulted in territorial coverage defects of class IV da neurons without altering microtubule stability (Figures 1 and 3), suggesting that *shot* may be serving pro-maintenance roles in these highly plastic dendrite arbors. One positive regulator of dendrite maintenance is non-centrosomal microtubule-organizing centers (ncMTOCs) such as Golgi outposts and endosomal Wnt signaling proteins at dendritic branchpoints (Ori-McKenney et al., 2012; Zhou et al., 2014; Yalgin et al., 2015; Harterink et al., 2018; Weiner et al., 2018, 2020; Liang et al., 2020; Mukherjee et al., 2020; Wilkes and Moore, 2020; Yang and Wildonger, 2020; Yagoubat and Conduit, 2026). For example, Golgi outposts co-localize with sites of EB1::GFP comet formation in dendritic branchpoints of class IV da neurons, and Golgi vesicles can promote microtubule nucleation *in vitro* (Ori-McKenney et al., 2012). The minus-end microtubule nucleator *Patronin* (CAMSAP1 in mammals) is required for proper dendritic microtubule polarity in da neurons, and *Patronin* is recruited by *shot* in *Drosophila* follicle cells and oocytes to regulate microtubule polarity at ncMTOCs (Khanal et al., 2016; Nashchekin et al., 2016; Feng et al., 2019). As our data suggested *shot* maintains territorial coverage in class IV da neurons and GFP-tagged Shot localized to dendritic branchpoints, we propose that *shot* may recruit *Patronin* for minus end-mediated microtubule growth in these neurons. Consistent with this, Davies et al. (2025) demonstrate that *shot* is required for dendritic microtubule orientation in class IV da neurons (Davies et al., 2025). Collectively, whether shot acts as a regulator of dendritic ncMTOCs, perhaps by recruiting Patronin to dendritic branchpoints, is an important open question with implications for understanding how minus-end microtubule dynamics contribute to arbor maintenance.

Alternatively, reduced territorial coverage of class IV da neurons with *shot* knockdown may have been an indirect result of *shot’s* inhibition of DLK/JNK signaling, which exerts bimodal control of dendritic and axonal growth. That is, *shot* knockdown activates DLK/JNK signaling that directs neurons to preferentially grow axons while concomitantly suppressing dendritic growth (Valakh et al., 2013; Wang et al., 2013). *Shot* loss-induced defects in territorial coverage of plastic class IV da neurons were DLK-dependent but JNK-independent, as evidenced by unchanged Puc::GFP levels (Figure 4). Strikingly, however, *Wallenda* overexpression in class IV da neurons also failed to increase Puc::GFP levels despite drastically reducing territorial coverage (Figure 5). This was at odds with established literature showing that homozygous mutants for *highwire,* an upstream E3 ubiquitin ligase that negatively regulates *Wallenda* protein levels and activity, increased Puc::LacZ levels in class IV da neurons (Wang et al., 2013). Additionally, increased complexity of motor neuron terminals in *highwire* mutants is phenocopied by *Wallenda* overexpression, and *Wallenda* overexpression is sufficient to increase Puc::LacZ in motor neurons (Collins et al., 2006; Xiong et al., 2010). Taken together, these findings suggest that plastic class IV da neurons may increase *highwire* activity or expression as a result of *shot* loss to keep DLK/JNK signaling low. Alternatively, non-canonical DLK signaling, perhaps through nemo kinase and its interactions with Hippo signaling (Emoto et al., 2006; Parrish et al., 2007; Merino et al., 2009), may underlie JNK-independent *shot* loss phenotypes.

### Identification and Functional Implications of Novel Shot Foci in Dendrites

Across both neuronal and non-neuronal cells, expression of GFP-tagged *shot* transgenes results in diffuse localization, accumulation of Shot in a rod-like manner, or a mix of both. These localization patterns are broadly consistent with *shot’s* functions in promoting microtubule stability and microtubule growth. For example, GFP-tagged Shot in tracheal cells is localized at cell junctions, microtubule bundles, and diffusely throughout the cytoplasm (Lee and Kolodziej, 2002b; Lee et al., 2003; Ricolo and Araujo, 2020). Shot is also highly enriched at the apical and basal surfaces of epidermal muscle attachment cells, accumulating at the termini of prominent microtubule bundles (Gregory and Brown, 1998; Subramanian et al., 2003; Bottenberg et al., 2009; Alves-Silva et al., 2012). In neuronal axons, Shot accumulates in a rod-like manner to axonal growth cones and along axonal shafts (Sanchez-Soriano et al., 2009). In *Drosophila* oocytes, however, Shot accumulates alongside Patronin as individual foci (Nashchekin et al., 2016), similar to what we observed in the dendrites of class I da neurons (Figure 2).

We speculated above that *shot* may serve minus-end mediated microtubule growth functions in class IV da neurons via these interactions with Patronin, but this is not consistent with *shot* loss increasing dendrite branch number in class I da neurons (Figure 1d-e). In addition to *Patronin’s* roles in minus end-mediated dendritic microtubule growth, it also functions in dendritic outgrowth and stabilization of microtubule minus ends in da neurons and cultured mammalian neurons (Goodwin and Vale, 2010; Hendershott and Vale, 2014; Jiang et al., 2014; Yau et al., 2014; Feng et al., 2019). Therefore, we propose that Shot foci work to stabilize microtubule minus ends and inhibit excessive minus-end mediated microtubule growth in neuronal dendrites, possibly alongside *Patronin.* While the excessive dendrite growth we observed in class I da neurons with *shot* loss was DLK-dependent and upregulated JNK signaling, *Wallenda* overexpression, which also increased JNK signaling in class I neurons, failed to produce excessive dendrite growth. This indicates that excessive dendrite growth is not simply a consequence of JNK activity in class I da neurons, and points to specific functional roles for Shot foci in the dendritic arbors of stable neurons. Dissecting the mechanistic underpinnings of these Shot foci in stable dendrites could yield novel insights into mechanisms that restrict excessive dendrite growth.

### Shot as a Key Regulator of Dendrite Regeneration

Several studies have established the role of *shot* in developing neurons, but fewer have examined how *shot* regulates the response to injury. After axon injury, neurons with depleted *shot* levels initiate regenerative programs more rapidly, likely through elevated DLK/JNK signaling priming neurons for axonal regrowth (Valakh et al., 2013, 2015). Despite opposing roles for *shot* in maintaining the dendrite arbors of class I da and class IV da neurons, we find that *shot* promotes dendrite regeneration in both cell types. This is likely due to *shot’s* function in MT extension, as *shot* binds the microtubule plus end-interacting protein EB1 to guide microtubule growth and regenerated dendrites display plus end-out microtubule polarity (Alves-Silva et al., 2012; Song et al., 2012; Hahn et al., 2021). However, *shot* may also be promoting dendrite regeneration by recruiting *Patronin*, or regulating the activity of ncMTOCs. This is supported by evidence that *Patronin* inhibits dendrite regeneration in class IV da neurons (Feng et al., 2019). Furthermore, while full-length Shot accumulates at dendrite tips in a rod-like manner, Shot puncta are visible throughout the regenerated dendrite arbor (Figure 7), suggesting possible resumption of its maintenance functions as in uninjured dendrites (Figure 2).

By expressing a neuronal Shot.LA isoform or a non-neuronal Shot.LC isoform in class I da neurons, we provide evidence that both actin-binding domains of Shot are required for its proper localization in regenerated dendrites. These findings are consistent with the requirement of these domains in *de novo* dendrite and axon growth (Bottenberg et al., 2009). We further assessed accumulation patterns of Shot domain mutants missing either the plakin, rod, calcium-binding, or microtubule-binding domain. These Shot domain mutants have been expressed in embryonic motor neurons situated in the *Drosophila* ventral nerve cord, but assessment of their subcellular distribution in dendrites was complicated by the compact nature of dendritic arbors in the CNS environment (Bottenberg et al., 2009). Conversely, expression of these Shot domain mutants in da neurons allowed us to robustly assess their accumulation patterns in regenerated dendrites. All domain mutants failed to phenocopy full-length Shot accumulation numbers, primarily due to their inability to reform small punctate accumulations. Notably, the plakin and rod domains were dispensable for rod-like accumulation, consistent with their non-essential role in dendrite development (Bottenberg et al., 2009). However, deeper assessment of their localization in regenerated dendrites revealed distinct functional roles for the plakin and rod domains. Taken together, these results suggest that all domains of Shot are required for the resumption of Shot foci, Shot’s plakin domain is dispensable for its microtubule growth-promoting functions, and Shot’s rod domain is essential for trafficking out to regenerating dendrite tips.

In summary, we find opposing roles for *shot* in regulating dendrite maintenance across morphologically and cytoskeletally distinct neuron types. These opposing roles in dendrite maintenance coincide with distinct accumulation patterns of Shot, in addition to cell type-specific differences in *shot* loss-induced DLK signaling. These results contribute to our current understanding of the role of *shot* in dendrite maintenance by demonstrating that *shot* can restrict dendrite growth in a context-dependent manner. Furthermore, a previously undescribed accumulation of Shot into foci distributed across stable dendrite arbors reveals a new mode of how Shot can accumulate in neurons. Our work also deepens our understanding of *Wallenda*/DLK signaling in neurons by suggesting that some neurons may be more resilient to *Wallenda*/DLK signaling activity. Lastly, we advance our understanding of dendrite regeneration by implicating *shot* as a key regulator of this biological process across neuron types, regardless of its opposing roles in dendrite maintenance. Future studies should decipher how dendritic Shot foci restrict excess dendrite growth and dissect the role of *shot* in regulating ncMTOC activity. Together, these findings establish a framework for investigating the context-dependent roles of *shot* in neuronal dendrites.

## Materials and Methods

### Drosophila Husbandry and Genetics

#### Husbandry

Adult flies were reared and maintained at room temperature (RT, ∼22.5C). Eggs were collected by transferring adult flies from each cross into a bottle capped with a grape juice agar plate and a spot of yeast paste. Scratches were made in the grape juice agar to encourage egg laying. Egg laying occurred over 2-4 hours, depending on animal numbers in crosses. Experimental larvae were reared and maintained in a humidified RT 22.5°C incubator until the desired developmental timepoint.

#### Genetics

All genotypes for individual figure panels can be found in the supplementary material. Most fly stocks were obtained from the Bloomington *Drosophila* Stock Center (BDSC), unless otherwise noted. Stock identifiers are listed as BDSC #. The following fly lines were used: ppk-Gal4 (BDSC #32079), 2.21-Gal4 (Grueber et al., 2003), UAS-CD4::tdTomato (BDSC #35841 [chromosome II] and #35837 [chromosome III]), 109(2)80-Gal4, UAS-mCD8::GFP (BDSC #8768) (Gao et al., 1999; Lee and Luo, 1999), ppk-CD4::tdTomato (BDSC # 35845), P{Wee}tau^304^ (Tau::GFP in the text, courtesy of Melissa Rolls), Puc::GFP (courtesy of Catherine Collins), ppk-Gal4, ppk-CD4::tdGFP, ppk-eGFP, nls-BFP-UAS-Dicer2 (courtesy of Melissa Rolls, ppk-CD4::tdGFP generated in (Han et al., 2011)), UAS-GMA (BDSC #31776), UAS-*Wallenda* (BDSC #51642). The following RNAi lines were generated by the Transgenic RNAi Project at Harvard Medical School (Perkins et al., 2015): UAS-shot.RNAi^HMJ23381^ (BDSC #64041), UAS-shot.RNAi^GLO1286^ (BDSC # 41858, *shot*.RNAi (B) in figures), UAS-yTub37C.RNAi (BDSC #32513), UAS-luciferase.RNAi (BDSC # 31603), UAS-*Wallenda*.RNAi (BDSC #27525). The following fly lines were generated by Seungbok Lee and Peter Kolodziej (Lee and Kolodziej, 2002a) and used in this study to visualize Shot in da neurons: UAS-Shot.LA::GFP (BDSC # 29044), UAS-Shot(LC)::GFP (BDSC #29042), UAS-Shot^ΔRod1^::GFP (BDSC #29040), UAS-Shot^ΔPlakin^::GFP, *shot*^3^ (BDSC #29649), UAS-Shot^ΔEF^::GFP (BDSC #29039), UAS-Shot^ΔGAS2^::GFP (BDSC #29041).

### Imaging Larval da Neurons

To immobilize larvae for imaging, a 4% agarose pad was created between two glass slides. Once solidified, the pad was cut into a square and reoriented into a diamond configuration. A small amount of vacuum grease was placed on each side of the agarose pad and a drop of glycerol was placed on the pad itself. A larva was picked from the egg collection plate, rinsed in water, then placed into a 35mm cell culture dish containing a piece of cotton soaked with isoflurane for anesthetization until cessation of mouth hook movement. Anesthetic was not used for larvae aged 96 h AEL or younger because these animals were sufficiently immobilized without isoflurane exposure. The larva was then placed onto the glycerol mounting medium, and a coverslip was gently pressed onto the vacuum grease to align the larva. Two pieces of tape were then applied perpendicular to the long axis of the slide on either side of the coverslip to fully immobilize the larva. Z-stacks (0.5µm-1µm slice intervals) of sensory neurons in larval segments A2-A5 were then acquired using a Zeiss LSM980, Zeiss LSM900, Zeiss LSM700, or Leica SP8 confocal microscope. The same microscope model was used for all images acquired within a given dataset. A 488 nm laser was used to image GFP fluorophores and a 561 nm laser was used to image tdTomato fluorophores. A 20x/0.8 NA dry objective was used to image whole dendrite arbors for morphological analyses. A 40x/1.3 NA water immersion Plan-Apochromat objective was used to image Shot localization in uninjured neurons and regenerated dendrite arbors, in addition to conducting laser injuries. A 63x/1.4 NA oil immersion Plan-Apochromat objective was used to image Puc::GFP levels.

### Injuring Larval da Neurons

Larvae at 96 h AEL were immobilized as described above, without anesthetic. A tunable near-infrared Ti:Sapphire laser was used for two-photon excitation-mediated laser injury on a Zeiss LSM980 confocal microscope. Injuries were performed to all primary dendrites of neurons in segments A2-A5 using a 12 x 12 pixel square ROI (∼1 µm^2^) completely surrounding primary dendrite branches. The laser wavelength was set to 860 nm, and a laser power of 100% (∼1570 mW) was delivered for 75 iterations in the 12 x 12 pixel ROI for ∼1 second. A 40x/1.3 NA water immersion Plan-Apochromat objective was used to image and injure neurons. After injury, larvae were returned to larval rearing plates with grape juice agar. Neurons were re-imaged for assessment of dendrite regeneration 48 hours post-injury as described above in “Imaging Larval da Neurons.”

### Imaging v’ada Neurons in the Adult Drosophila Abdomen

Male flies were first immobilized in an empty 100 mL plastic vial plugged with a cotton ball soaked in isoflurane. Using an acrylic plastic disc as previously described (DeVault et al., 2018), adult flies were mounted on their left side, and the wings and legs were carefully repositioned to provide clear visualization of the v’ada neurons. A small drop of glycerol was added to cover only the abdomen. Small magnetic strips secured a 24 x 50 mm coverslip (VWR) on top of the immobilized fly. The v’ada neurons were imaged using a 20x/0.8 NA dry objective on a Zeiss LSM980 microscope.

### Immunostaining of Larval Fillet Preparations

To immunostain for Shot in larval fillet preparations, we followed previously published protocols (Tenenbaum and Gavis, 2016; Xu et al., 2024). Third-instar larvae were dissected in 35 mm dishes using insect pins, and body wall muscles were gently removed. All subsequent steps were carried out on a shaker set to 60 rpm in a light-protective container to minimize photobleaching of genetically-driven CD4::tdTomato. Fillet preparations were fixed for 25 minutes in 4% paraformaldehyde, washed three times, 10 minutes each in 0.1% Triton X-100 in PBS (PBST), then transferred to microcentrifuge tubes in 5% normal goat serum (NGS). Samples were blocked for 20 minutes at RT, and incubated overnight at 4°C with a mouse anti-Shot mAbRod1 antibody that recognizes the rod domains of the Shot.LA isoform (1:50 dilution, DSHB, antibody registry RRID:AB_528467). Samples were washed six times, 10 minutes each with PBST at RT, incubated for 2 hours with Alexa Fluor 488 goat anti-mouse secondary antibody (Invitrogen A10680, 1:750) in 5% NGS, then washed six times, 10 minutes each with PBST at RT. Fillet preparations were mounted onto slides; the heads and tails of animals were cut off to achieve a flat body wall preparation. Approximately 50 µL of VectaShield Antifade Mounting Medium (Vector Laboratories) was placed on the fillets before placing the coverslip, and samples were immediately imaged using a 20x/0.8 NA Plan-Apochromat objective on a Zeiss LSM980 microscope.

### Quantifications

All data were quantified using Fiji/ImageJ. Quantifications are described in the order by which they appear throughout all figures. All data were analyzed by researchers blind to animal genotype.

#### Territorial Coverage of Larval Uninjured class IV ddaC Neurons

To quantify territorial coverage of class IV ddaC neurons, we used a modified version of the “Internal Coverage” macro (Sears and Broihier, 2016). This macro overlays a grid of squares on images of class IV ddaC neurons, and then analyzes territorial coverage as the proportion of squares with positively-filled dendrites. We modified the macro to fill positive squares with respective genotype colors and automated the macro to iteratively quantify positively-filled squares for all images in a given dataset. Images of class IV ddaC neurons were median filtered, maximum-intensity projected, Gaussian blurred, and subjected to auto local thresholding using the Phansalkar method, and background particles were removed using the “Analyze Particles” function. The exact values used for each function differed depending on the fluorescent reporter used but were identical between genotypes within a dataset. Generally, images were median filtered with a radius of 1.0, Gaussian blurred with a radius of 1.0, subjected to auto local thresholding with a radius of 30, and background particles smaller than 100 pixels were removed. The proportion of positively-filled squares (i.e. those with dendrites) was used to quantify differences in territory coverage between genotypes.

#### Branch Number and Total Dendrite Length of class I ddaE Neurons

To quantify dendrite branch number and total dendrite length (µm) in class I ddaE neurons, we used the Neuroanatomy plugin Simple Neurite Tracer (SNT) software (Arshadi et al., 2021). SNT enables semi-automated tracing of dendritic processes, designated as “paths” within the software. Paths were saved as .traces files and path properties were exported as .csv files to obtain path counts (branch number) and path lengths (total dendrite length).

#### Anti-Shot Immunostaining

Images were quantified by first creating maximum intensity projections, then manually drawing ROIs corresponding to class I ddaE neuron somas (white arrowheads in Figure S1a). Mean anti-Shot intensity values were measured and plotted.

#### Territorial Coverage of Uninjured Adult v’ada Neurons

To quantify territorial coverage of adult class IV v’ada neurons, images were first processed as described for territorial coverage of larval class IV ddaC neurons. As mentioned in the text, our ppk-Gal4, UAS-CD4::tdTomato labeling approach resulted in both class IV v’ada neurons (purple arrows to soma) and class II neurons (red arrows to soma) expressing CD4::tdTomato (Figure S1h). We therefore restricted ROIs to the region near the ventral midline, as this area is enriched for class IV v’ada dendrites and receives minimal contribution from class II neuron dendrites (Shimono et al., 2009). Territorial coverage analysis was restricted to these ROIs, and the proportion of positively-filled squares was plotted.

#### Morphological Analysis of Shot Accumulations

Images were processed in a similar manner to those subjected to territorial coverage analysis of class IV ddaC neurons (median filtered, maximum-intensity projected, gaussian blurred, auto local thresholded using the Phansalkar method, and background-particle filtered using the “Analyze Particles” function). Dendrite masks of Shot puncta were created by exporting SNT-generated paths to the ROI manager. Paths were enlarged to width = 20 pixels (width = 10 pixels for puncta in regenerated dendrites), and a single ROI with all paths was created by filling a blank image canvas. Any remaining signal including non-specific background, soma signal, axon signal, and signal from adjacent neurons, was removed using the “Clear Outside” function on the main image. Shape descriptors were selected in “Set Measurements” to quantify aspect ratio and size. The dendritic Shot mask was then analyzed using the “Analyze Particles” function, with “Display Results” selected for frequency distributions of individual Shot accumulations and “Summarize” selected for per neuron measures of Shot accumulations. For binning, percentile thresholds were derived from Shot accumulations within the Shot.LA group.

#### Tau::GFP and CD4::tdTomato Levels

Images were maximum-intensity projected. We used the segmented line tool (width = 2 pixels, spline fit) to draw ROIs corresponding to neurites and the polygon selection tool to draw ROIs corresponding to somas. Measures were obtained for all ROIs in both channels, and mean intensity was plotted. For class IV ddaC somas, median intensity was used as CD4::tdTomato frequently accumulated in the soma, resulting in signal bleed-through into the green channel and pixel saturation.

#### Puc::GFP Levels

Circular ROIs were created in da neuron nuclei using the oval selection tool. We chose the z-slice with the highest intensity Puc::GFP signal for quantification, as previously described (Stone et al., 2014). Mean Puc::GFP intensity was plotted.

#### Regenerated Area of Injured class IV ddaC neurons

Images were maximum-intensity projected, and the polygon selection tool was used to draw ROIs around regenerated class IV ddaC dendrite arbors. The area was measured and plotted.

#### Sholl Analysis

To quantify accumulation complexity in mutants, Sholl analysis was performed using the automated Sholl plugin integrated into SNT (Ferreira et al., 2014). The center of the analytical shell grid was anchored at the midpoint of the neuronal soma. Concentric circles were defined starting at a minimum radius of 10 µm from the soma center, increasing step-wise at 1 µm increments to a maximum radius of 388 µm. The number of accumulation intersections per sphere was recorded to map Shot^ΔPlakin^ and Shot^ΔRod1^ distribution as a function of distance from the soma.

### Statistical Analyses

All statistical tests were performed in GraphPad Prism software, and statistical significance was defined as p < 0.05. Specific statistical tests used for each dataset can be found in figure legends. Generally, a t-test was used to compare two groups and a one-way ANOVA with multiple comparisons correction was used to compare multiple groups.

## Acknowledgements

We thank Adeela Syed, PhD and the Optical Biology Center at UCI for extensive use of their microscopes. This study was made possible in part through access to the Optical Biology Core Facility of the Developmental Biology Center, a shared resource supported by the Cancer Center Support Grant (CA-62203) and NIH-S10OD032327-01. Stocks obtained from the Bloomington Drosophila Stock Center (NIH P40OD018537) were used in this study. KTP is a fellow of the Hellman Foundation and the Rose Hills Foundation. This work was supported by startup funds from the UC Irvine Dunlop School of Biological Sciences (to KTP) and NIH grant R00NS097627 (to KTP). VND was supported by the UCI BioSci Postdoc excellence fellowship. RB was funded by the California Institute for Regenerative Medicine Predoctoral Research Training Grant (Award # EDUC4-12822).

## Author contributions

Conceptualization: VND and KTP. Methodology: VND. Investigation: VND, RB, and VN. Formal Analysis: VND and KTP. Visualization: VND. Writing – Original Draft: VND. Writing – Review & Editing: VND and KTP. Supervision: KTP. Funding Acquisition: KTP.

