## Supplementary Figures and legends for "Opposing Roles for the Spectraplakin Short Stop in Stable and Dynamic Dendrites Reveal Divergent DLK Signaling and a Role in Dendrite Regeneration"

Duarte_FigS1

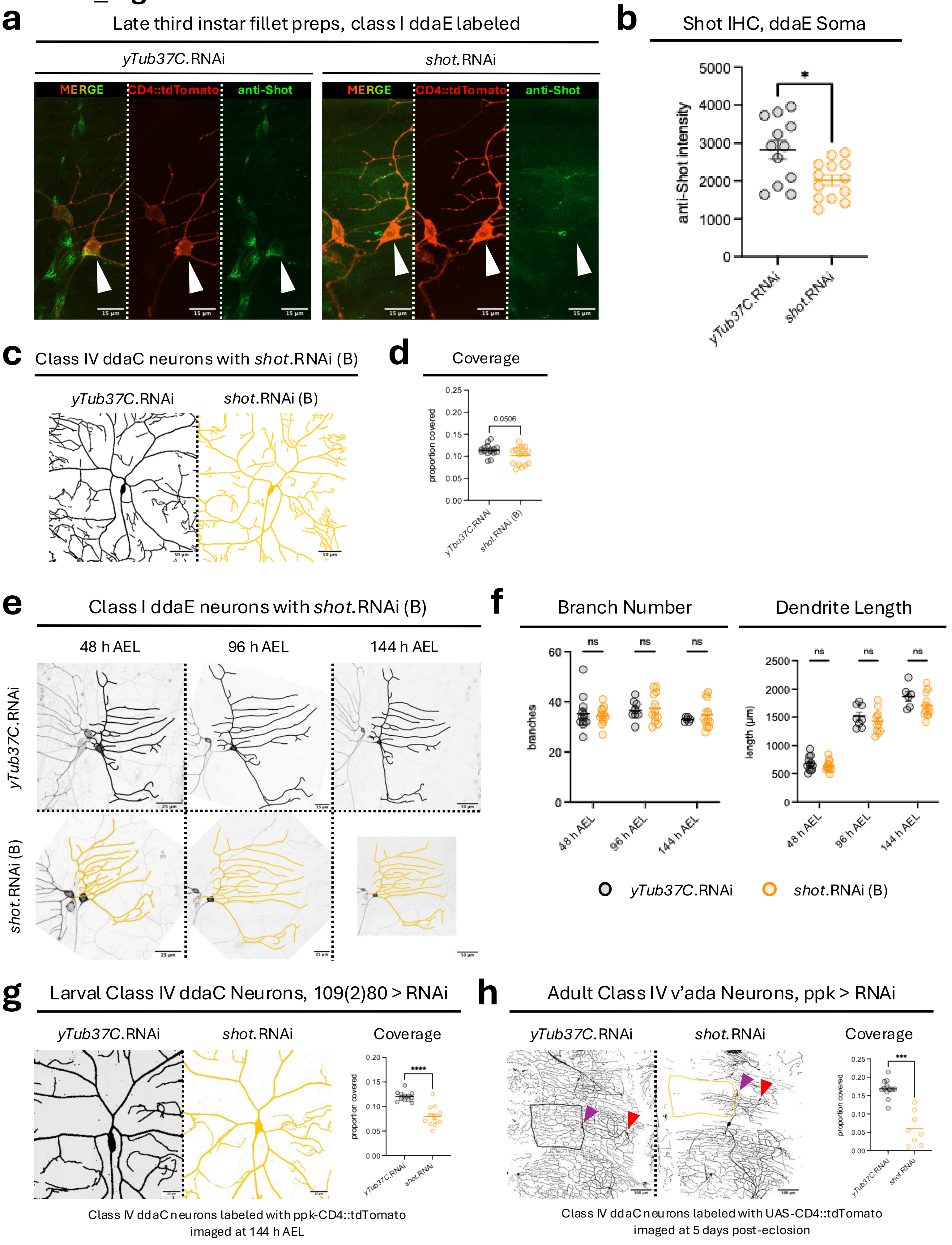

### Duarte_FigS1

**Figure S1**: **(a)** Immunohistochemical staining of anti-Shot in late third-instar fillet preparations with control *yTub37C* knockdown or *shot* knockdown. Class I ddaE neurons are genetically labeled using 2.21-Gal4 and UAS-CD4::tdTomato. Arrowheads point to class I ddaE somas. **(b)** Mean anti-Shot intensity values. Welch's t-test. p = 0.0108. N = 12-13 cells/group, 3 larvae/group, 3-6 cells/larvae. **(c)** Class IV ddaC neurons in 144 h AEL larvae with control *yTub37C* knockdown or *shot* (B) knockdown. **(d)** Territory coverage showed a trending decrease in neurons with *shot* (B) knockdown. Welch's t-test. p = 0.0506. N = 16 cells/group, 4-5 larvae/group, 2-4 cells/larvae. **(e)** Repeatedly imaged class I ddaE neurons in 48/96/144 h AEL larvae with control *yTub37C* knockdown or *shot* (B) knockdown. **(f)** Branch number and total dendrite length showed no changes in neurons with *shot* (B) knockdown. Multiple unpaired t-tests with Welch correction and Holm-Šídák multiple comparisons correction. N = 6-12 cells/group, 3-6 larvae/group, 2 cells/larvae. **(g)** Class IV ddaC neurons in 144 h AEL larvae with control *yTub37C* knockdown or *shot* knockdown driven by 109(2)80-Gal4. Class IV ddaC neurons are labeled using ppk-CD4::tdTomato. Territory coverage showed a significant decrease in neurons with *shot* knockdown. Welch's t-test. N = 12 cells/group, 4 larvae/group, 3 cells/larvae. **(h)** Class IV v'ada neurons in the adult abdomen with control *yTub37C* knockdown or *shot* knockdown driven by ppk-Gal4. Class IV v'ada neurons are labeled using UAS-CD4::tdTomato. Territory coverage showed a significant decrease in neurons with *shot* knockdown. Welch's t-test. N = 8-12 cells/group, 8-12 adults/group, 1 cell/adult. Mean ± SEM is plotted for all graphs.

### Duarte_FigS3

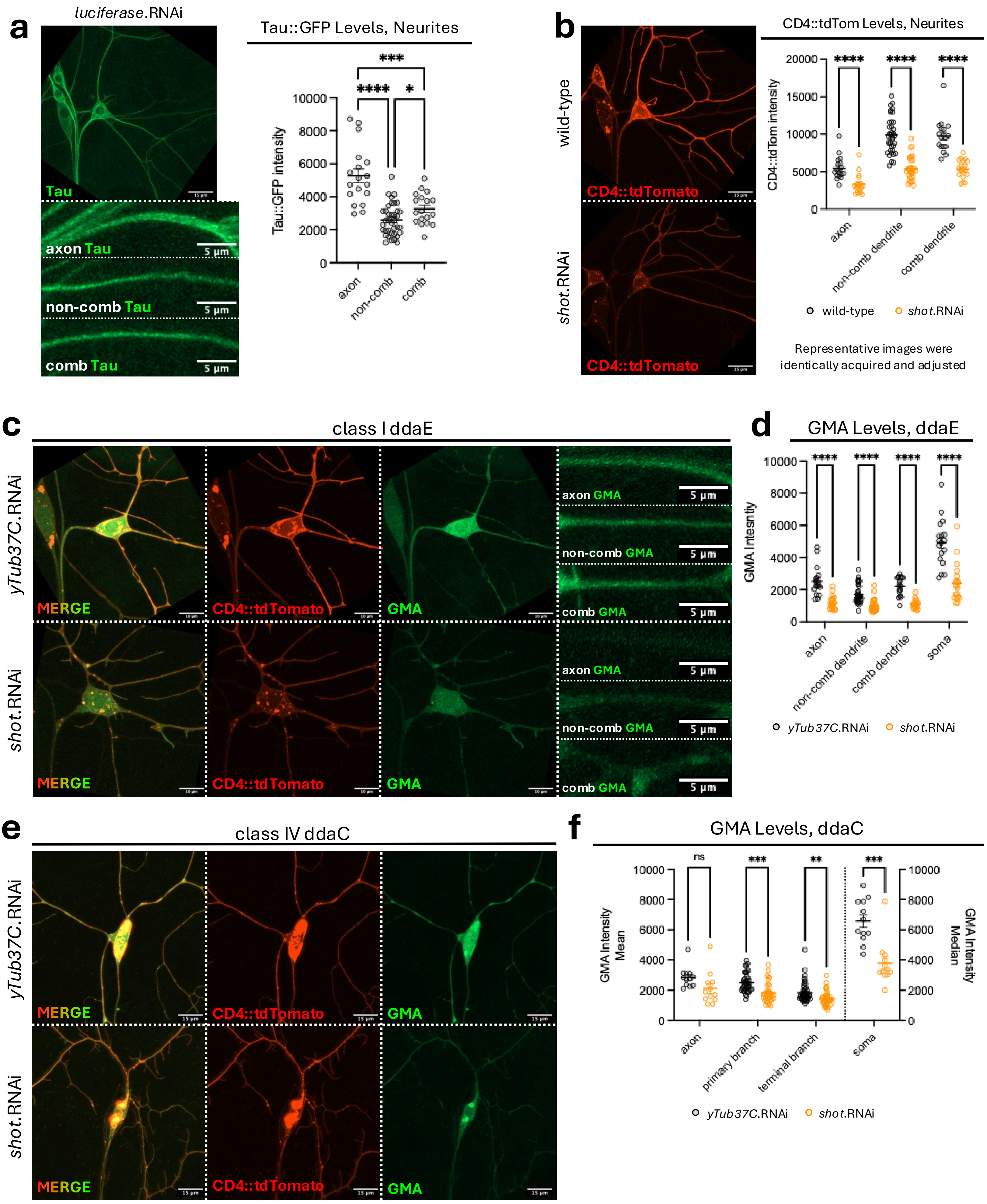

### Duarte_FigS3

**Figure S3: (a)** Tau::GFP in 144 h AEL larvae in wild-type class I ddaE neurons, with zoom-ins for the axon, primary non-comb dendrite, and primary comb dendrite. Tau::GFP levels across neurite compartments show significant differences between all structures. Brown-Forsythe and Welch ANOVA test with Dunnett's T3 multiple comparisons correction. N = 18 cells/neurite, 6 larvae/neurite, 3 cells/larvae, 18 axons, and 37 non-comb dendrites, and 18 comb dendrites. **(b)** Class I ddaE CD4::tdTomato levels in 144 h AEL larvae in wild-type or *shot* knockdown neurons. CD4::tdTomato levels across neurite compartments show significant decreases in *shot* knockdown neurons across all structures. Multiple unpaired t-tests with Welch correction with Holm-Šídák multiple comparisons correction. N = 18 cells/genotype, 6 larvae/genotype, 3 cells/larvae, 18 axons/genotype, 18 comb dendrites/genotype, and 30-37 non-comb dendrites/genotype. **(c)** Class I ddaE neurons in 144 h AEL larvae with control *yTub37C* knockdown or *shot* knockdown. Neurons are labeled using 2.21-Gal4 and UAS-CD4::tdTomato. Stable F-actin is labeled using UAS-GMA. Zoom-ins show GMA in the axon, primary non-comb dendrite, and primary comb dendrite. **(d)** GMA levels showed a significant decrease in axons, primary non-comb dendrites, primary comb dendrites, and in the soma in neurons with *shot* knockdown. Multiple unpaired t-tests with Welch correction with Holm-Šídák multiple comparisons correction. N = 18 cells/genotype, 6 larvae/genotype, 3 cells/larvae, 18 axons/genotype, 31-35 non-comb dendrites/genotype, 18 comb dendrites/genotype, and 18 soma/genotype. **(e)** Class IV ddaC neurons in 144 h AEL larvae with control *yTub37C* knockdown or *shot* knockdown. Neurons are labeled using ppk-Gal4 and UAS-CD4::tdTomato. Stable F-actin is labeled using UAS-GMA. **(f)** GMA levels showed no change in axons, but a significant decrease in primary branches, terminal branches, and in the soma in neurons with *shot* knockdown. Median intensity was used for soma measures as CD4::tdTomato frequently accumulated in this compartment, resulting in signal bleed-through into the green channel and pixel saturation. Multiple unpaired t-tests with Welch correction with Holm-Šídák multiple comparisons correction. N = 12 cells/genotype, 4 larvae/genotype, 3 cells/larvae, 12 axons/genotype, 36-40 primary branches/genotype, 41-47 terminal branches/genotype, and 12 soma/genotype. Mean ± SEM is plotted for all graphs.

Duarte_FigS6

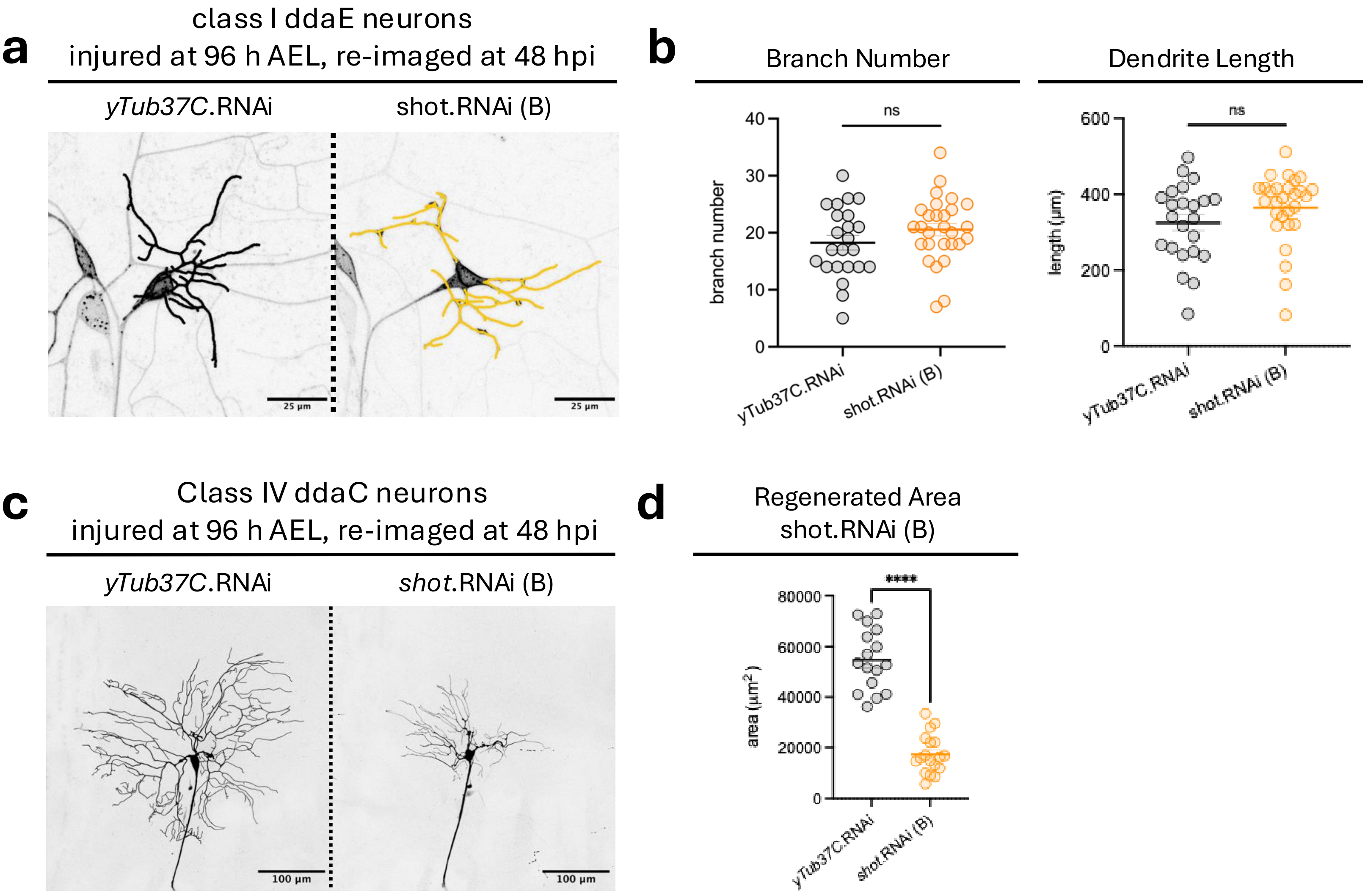


**Figure S6: (a)** Class I ddaE neurons at 48 hpi in larvae with control *yTub37C* knockdown or *shot* (B) knockdown. Injuries occurred 96 h AEL. Traces of regenerated dendrites are overlaid. **(b)** Branch number and total dendrite length showed no changes in neurons with *shot* (B) knockdown. Welch's t-test. N = 23-28 cells/genotype, 9 larvae/genotype, 1-3 cells/larvae. **(c)** Class IV ddaC neurons at 48 hpi in larvae with control *yTub37C* knockdown or *shot* (B) knockdown. Injuries occurred 96 h AEL. **(d)** Regenerated area showed significant decreases in neurons with *shot* (B) knockdown. Welch's t-test. N = 23-28 cells/genotype, 9 larvae/genotype, 1-3 cells/larvae. Mean ± SEM is plotted for all graphs.

### Duarte_FigS7

**a**

Class IV ddaC neurons
injured at 96 h AEL, re-imaged at 48 hpi

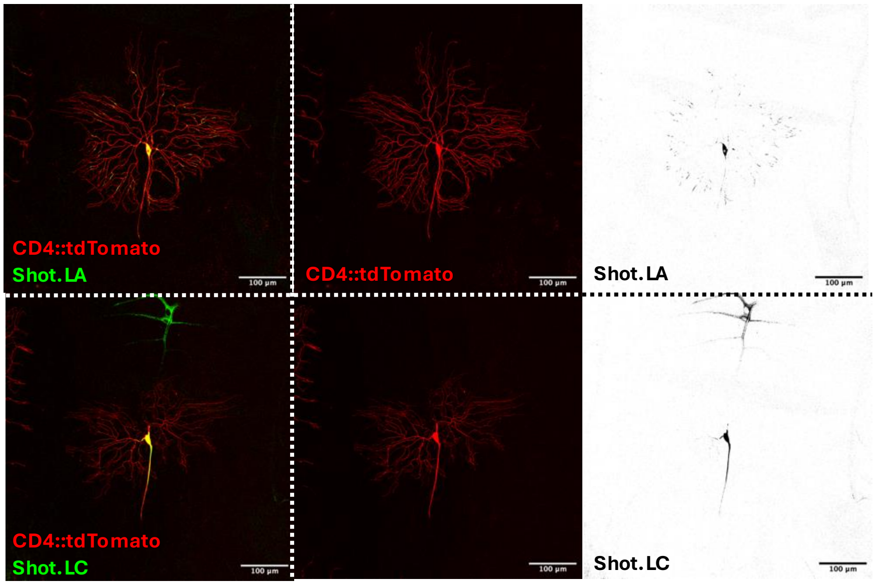


**b**

Sholl Analysis,
Dendritic Shot$^{\Delta Plakin}$ and Shot$^{\Delta Rod1}$ Masks

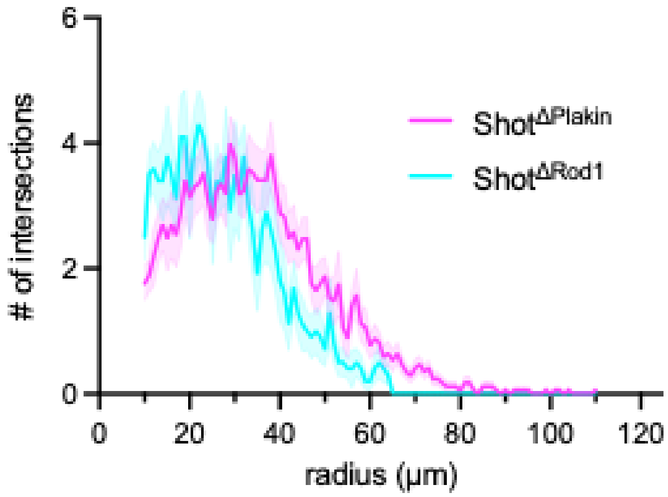


**Figure S7: (a)** Class IV ddaC neurons at 48 hpi in larvae overexpressing GFP-tagged neuronal Shot.LA or non-neuronal Shot.LC. Injuries occurred 96 h AEL. **(b)** Sholl analysis of dendritic Shot$^{\Delta Plakin}$ and Shot$^{\Delta Rod1}$ masks with a starting radius of 10 µm from the soma and a step size of 1 µm.
